# Assessing the potential of ancient protein sequences in the study of hominid evolution

**DOI:** 10.1101/2025.04.08.647730

**Authors:** Ioannis Patramanis, Laurits Skov, Enrico Cappellini, Fernando Racimo

**Author notes:** November 10, 2025.

## Abstract

Palaeoproteomic data can provide invaluable insights into hominid evolution over long timescales. Yet, the potential and limitations of ancient protein sequences to resolve evolutionary relations between species remains largely unexplored. In this study, we aim to quantify how much information about these relations can be obtained from limited ancient protein data, at the scale that is currently available or will be available in the near future. We harness sequence alignments of 12 enamel and collagen proteins that have been previously reported in fossil material that is at least 1 million years old. We utilise in silico translations of hominid DNA sequences of these proteins and highlight their differential sequence conservation, indicating some of them contain much larger amounts of information than others. We also evaluate the extent to which inferred topologies from protein data differ from inferred topologies from the more informationally-dense DNA data. We show that the former may sometimes lead to inferences of the wrong tree topology due to the informational loss that comes when working with peptide data. Additionally, we determine the number of concatenated proteins necessary to confidently reconstruct the population / species tree summarizing the relations between humans, chimpanzees and gorillas, as well as those between modern humans, Neanderthals and Denisovans. As expected, increasing the number of proteins in a concatenation enhances resolution, but we note that trees inferred from the full set of collagen and enamel proteins do not necessarily correspond to population trees inferred from genome-wide data. We show this is especially the case in the closely related groups of our recent ancestors. We further demonstrate that while a number of proteins fall within archaic introgressed haplotypes of present day humans, ancient admixture is not the main source of the observed tree incongruence. Our study underscores the potential and limitations of utilising palaeoproteomic data in deep time phylogenetic reconstructions, indicating that these will be aided not only by increased recovery of proteins in the future, but also by more careful modeling of evolutionary relations across the genome, beyond simply building single phylogenetic trees.

## Introduction

Understanding the evolutionary relationships between extinct humans and other hominin groups is a fundamental question in paleoanthropology. This problem has largely been approached via methodologies derived from comparative morphology, phylogenetics and, more recently, the study of ancient DNA [1, 2, 3, 4]. In the last decade, improvements in the extraction and sequencing of ancient peptides have provided researchers with yet another source of evidence to tackle this problem. Paleoproteomics is an emerging field that addresses the deep-time limitations of ancient DNA, which degrades faster than the peptides of specific proteins [5]. Thus, ancient peptide sequencing has enabled the study of evolutionary relationships between organisms that lived from tens of thousands [6] to millions of years in the past [7, 8]. It has also yielded valuable data in areas where DNA tends to be poorly preserved, due to humid and warm climates [9]. These include regions of the world such as southern Europe [10], southern Asia [11, 12] and Africa [13, 14], which are rich in archaic hominid fossil material. Palaeoproteomics thus holds the potential to explore questions that were previously impossible to address using morphological or DNA data alone, including resolving the species or population identity of hundreds of fragmentary fossil specimens for which limited or no DNA sequences are available. Although the study of ancient proteins holds great promise, it also harbors limitations, caused by the number of proteins that can be retrieved from ancient material, and by the nature of protein data.

While some studies have managed to recover tens or even hundreds of proteins from relatively young samples [15, 16, 17, 18, 19, 20, 21, 22, 23], only collagen type I, enamel-specific proteins and a few others have so far been retrieved from million years old mammalian fossil material [8, 11, 24, 25, 26]. As only a handful of protein sequences are so far recoverable, only small amount of useful genetic information can be obtained from them. The sequences of these proteins are also not complete; degradation over time breaks down the original proteins into smaller and smaller peptides. After thousands of years, many peptides do not survive this process and are therefore unrecoverable, while peptides that do, can incorporate ambiguous amino acids due to post-mortem chemical modifications [5].

Even when fully preserved, proteins inherently contain less phylogenetic information than DNA [27]. This is both due to the degenerate nature of the genetic code [28], and to the fact that negative selection tends to act upon protein coding sequences more strongly than in other regions of the genome, reducing sequence variation [29]. Consequently, ortholog proteins of closely related species often show limited or no variation. Amino acid mutations may also occur in the same position multiple times in different lineages, due to functional molecular constraints [30], leading to molecular convergence or homoplasy [31]. This convergence can create the illusion of close phylogenetic affinities and has been observed in multiple taxa [30, 32, 33, 34, 35, 36]. Furthermore, when only a few proteins sequences from a given species are available, it is difficult to establish whether observed variants are fixed or polymorphic within that taxon [12, 13]. All of the above reduce the utility of amino acid polymorphisms for reconstructing evolutionary relationships.

Moreover, small sets of ancient peptides can provide very limited information about the full ancestral recombination graph, i.e. the graph structure that describes the full genealogical relationships of a set of individual genomes [37, 38]. Reconstructions based on these peptides tend to focus on individual gene trees, which in turn provide only partial knowledge about overall population relationships. This may be because gene trees are affected by incomplete lineage sorting (ILS) in ancestral populations [39, 40], or because admixture events between populations may not be represented in such trees [41]. Both ILS and admixture are of concern in species or populations that are closely related to each other [42, 43, 44, 45, 46]. In African great apes, for example, multiple studies have detected both high levels of ILS (amounting to up to 30% percent of the genome) [40, 47, 48, 49], as well as possible past admixture episodes, in all 3 extant genera [50, 51, 52, 53].

Although most of these limitations have been previously acknowledged [10, 13, 54, 55], little quantitative work has been done to assess the potential and limitations of ancient protein data at resolving population relationships [56, 57, 58, 59]. In this study, we specifically focus on 12 collagen and enamel proteins and their ability to resolve evolutionary relationships between species. We focus on these proteins because they have been previously recovered from biological material older than a million years. We center our analysis on the Hominidae family and its well-studied genetic history, based on DNA evidence [60, 61, 62, 63], and leverage the high availability of genomic and protein data that exists for all four of its extant genera and some extinct populations [60, 64, 65, 66]. For each of the 12 proteins, we use only a single isoform, labeled as “canonical’ in Ensembl, as this is what has been previously recovered in ancient material. Lastly we investigate whether the recovery of peptides from a richer proteome, such as that of dentin or bone, would enhance evolutionary resolution (and if so, how much).

First, we measure the entropy and evolutionary conservation rates of these 12 proteins, using hominid alignments, ranking them based on the amount of information they provide and comparing them to other known conserved proteins. We further use the entropy metric to measure the informational loss that occurs when comparing intron-containing DNA alignments, to exon and to protein alignments of the same genetic locus. We additionally evaluate how inferred topologies differ between these alignments of different data type.

While ILS leads to topological mismatches between gene trees and the population tree, topological mismatches can also occur between the true and the inferred gene tree at a given locus, either when using DNA or protein sequences (see figure 1). DNA gene tree misinference can occur due to the inherent difficulties of reconstructing topologies from mutations, as well as from the information loss and errors that can take place during the sequencing process. Protein gene tree misinference, can occur due to the same reasons, but also the additional information loss that occurs in protein translation. Local tree reconstructions are also necessarily affected by difficulties in inferring recombination events and resulting topological changes along the genome. For simplicity in our empirical analysis, and given our focus on information loss in ancient peptides specifically, we assume that gene trees reconstructed from DNA sequences are accurate and use them as a proxy for the true gene trees. We also assume that the sequences we study are small enough to be characterized by a single gene tree. However, we note that both of these are strong assumptions that might not hold in reality.

**Figure 1:**
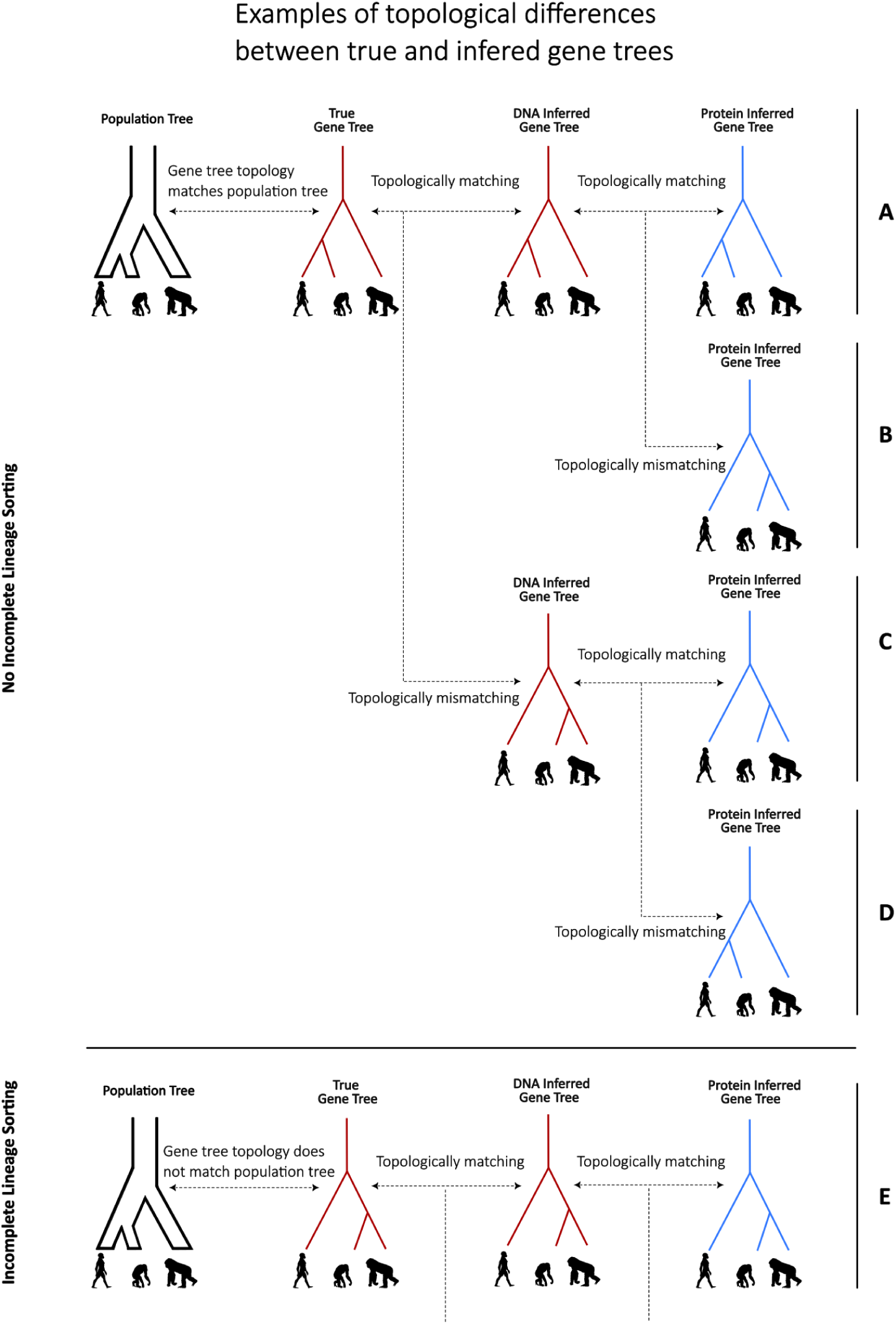
Schematic explaining different scenarios of tree topology discordance and concordance. The first column depicts the evolutionary relations between three different species (humans, gorillas and chimpanzees), as described by a simple population tree. The second column depicts the tree representing the genetic relation between 3 homologous segments at a specific locus of genomes obtained from these three species (a “gene tree”). The third column depicts a tree inferred using DNA sequence data, obtained from said homologous region. The fourth column depicts the gene tree inferred using protein data from the same region. We show a (limited) set of the many possible scenarios that can arise when comparing all the above trees with one another: **A.** No incomplete lineage sorting and all four trees agree with one another: the use of either DNA or protein data leads to the correct inference of the gene tree, which happens to agree with the population tree. **B.** No incomplete lineage sorting, but the protein gene tree differs from the true gene tree at the locus under study (e.g. due to the reduced information contained in peptide sequences, relative to DNA). **C.** No incomplete lineage sorting, but both the DNA- and protein-reconstructed gene trees are misinferred (e.g. due to very few genetic variants present at both the DNA and protein levels for correct tree resolution). **D.** No incomplete lineage sorting, but while the DNA-reconstructed gene tree is misinferred, the topology of the p_1_r_3_otein-reconstructed tree happens to match the true gene tree. **E.** There is actual incomplete lineage sorting (mismatch in topology between the true population tree and the true gene tree under study). In this specific case, the gene tree is also correctly inferred using either DNA or protein data. Other scenarios involving incomplete lineage sorting but misinferred topologies from either DNA data, protein data or both have been omitted for brevity. All silhouette images are reused from https://www.phylopic.org/.

We use an iterative analysis to compare protein-inferred phylogenetic trees with established tree topologies based on previously published genetic data. We estimate the number of combined proteins and amino acid variants required to reliably infer the population trees of 3 hominid genera and 3 hominin populations. We then repeat this iterative analysis by supplementing our 12 initial “deep time” proteins with 16 additional proteins, which have been experimentally recovered from bone or dentin [67, 68, 69], albeit so far, only from samples that are relatively younger.

## Materials and Methods

### Incomplete lineage sorting, DNA and proteins

We first selected 12 proteins that have previously been recovered from either tooth enamel (AHSG, ALB, AMBN, AMELX, AMELY, AMTN, COL17A1, ENAM, MMP20, ODAM) or bone material (COL1A1, COL1A2), from mammalian samples that are more than 1 million years old [8, 11, 24, 25, 26]. We then acquired the “canonical” isoform’s reference sequence for each of the proteins from Ensembl [70], for the following four hominid species: *Homo sapiens*, *Pan troglodytes*, *Gorilla gorilla*, *Pongo abelii*. We aligned the ortholog sequences from the four hominid species using Mafft [71] and reconstructed gene trees with PhyML [72], using each of the 12 ortholog alignments separately. We rooted the 12 generated trees using *Pongo abelii* as the outgroup and compared them to the population tree that best represents the relationships between those 4 species [61]. To compare our protein tree results, we repeated the same process but using the reference DNA sequences (combined exons and introns) of the genes corresponding to the 12 proteins instead of the amino acid sequences.

### Entropy and evolutionary conservation rates

To assess the conservation levels and the phylogenetic information of these 12 proteins, we calculated Shannon’s information theoretic entropy [73] and an evolutionary rate score [74] for each amino acid position on the sequence alignments. We used these two scores as approximations to the information content in these alignments, while respectively ignoring and accounting for the evolutionary distance between each sequence. We used Bio3d [75] for the entropy calculation and Rate4Site [74] for the evolutionary rate computation on the alignments of the four hominid species. We obtained a score for each position of each of the 12 protein alignments. We aggregated the metrics across each protein to obtain a total score, and also divided them by the length of the alignment, to obtain a sequence-wide average score. To account for within-species diversity, we used multiple individuals as representatives from each of the four species, using previously published translated proteomes] [76]. We randomly sampled 1 individual from each of the four species, repeated the calculations for 1000 repetitions and calculated the mean from all 1000 repetitions. To contextualize our results, we selected 5 proteins that have been previously reported as being either highly conserved or as containing hyper-variable segments [77] and included them in the entropy and evolutionary rate calculations. These included two highly conserved histones (H2BC3, H2BC9) and one ubiquitine (USP46) [78, 79], as well as two fibrinogen proteins (FGB, FGG) reportedly bearing a highly variable segment [77].

### Informational content: exons, introns and proteins

We assessed the differences in informational content between a DNA sequence containing both introns and exons, a version of the same sequence containing only exons, and a peptide version of the sequence (translated amino acids) for all 12 loci of our analysis. We applied Bio3d’s entropy scoring to all three data types of the same gene, for all 12 genes, and then ranked the results (see Sup. Material for details). We also divided each entropy metric by the length of the data type to compare the average information content per site, of each data type. Due to sequence length differences between DNA and protein data (with a 3 letter DNA codon corresponding to a single protein amino acid), we also applied a “length-correction” to this last measurement. In this correction we divided the entropy of each protein version of each gene by 3 (number of nucleotides that correspond to an amino acid), while keeping the introns-and-exons and exons-only version unaltered.

### Iterative phylogenetic analysis

We investigated how the concatenation of different numbers and different combinations of the 12 proteins might affect the topology of the inferred “consensus” tree, which is often taken as an estimate of the population or species tree. For this analysis we utilised a “hominid dataset”, consisting of *Homo sapiens*, *Pan troglodytes*, *Gorilla gorilla* and *Pongo abelii* (as an outgroup) and a second “hominin dataset” consisting of *Homo sapiens*, Neanderthals, Denisovans and *Pan troglodytes* (outgroup). To assess how the recovery of additional proteins, from different tissues affects phylogenetic analyses, we expanded the “hominin dataset”, creating a third “bone-dentin dataset”. This dataset consisted in the protein sequences that are most often recovered from dentin or bone tissue. In choosing which proteins to include in this anaysis, we utilised the list provided by Ruther et al. 2022 [67], which includes 20 proteins utilised in species identification: COL1A1, COL1A2, COL2A1, COL3A1, COL4A4, AHSG, COL5A2, ALB, BGN, COL5A3, COL5A1, CHAD, COL22A1, COL11A2, SERPINF1, F2, COL11A1, LUM, COL12A1, POSTN. Four of these 20 proteins (COL1A1, COL1A2, AHSG and ALB) were already included in the original 12 proteins, leading to a final combined dataset of 28 proteins.

In each iteration of this analysis, we carry out a concatenation using a subset of proteins sampled from the full set of proteins, reflecting the fact that not all proteins in the full set might be available in practice. The subset ranges in size from 1 (a single protein recovered) to all proteins recovered (either 12 or 28, depending on the tested dataset). One representative individual per population or species is randomly chosen and included in the alignment, as ancient protein studies are often limited to single individuals that are made to represent an entire species. For each concatenation, we build a phylogenetic tree and record the resulting topology. We then compare it to the underlying population tree, as inferred from past DNA studies. In total, we do this over 1,000 iterations per each number of proteins, sampling different sets of proteins and different representative individuals, in each turn.

We performed the same iterative analysis on each of the three datasets (“hominid”,“hominin”,“bone-dentin”). The analysis for the “hominid” and “hominin” datasets was repeated 1000 times for each *N* , with *N* ranging from 1 to 12, resulting in a total of 12,000 generated trees for each of the two datasets. The same process was applied to the bone-dentin dataset, with *N* ranging from 1 to 28, resulting in a total of 28,000 trees. For each iteration, we first picked *N* proteins, without replacement, out of the maximum number of proteins for that dataset. For each protein, one sequence from each of the four taxa was randomly selected from the samples available to us [80] and then the four orthologous sequences were aligned using Mafft. Each iteration (out of a thousand) thus generated a total of *N* protein alignments. The *N* alignments were then concatenated into a single alignment which was used to generate a phylogenetic tree using PhyML. The generated tree was also trimmed for very short and unsupported branches (see Sup. material), which were transformed into polytomies. The tree was then rooted using an outgroup taxon (*Pongo* for hominid set, *Pan* for hominin and bone-dentin set) and compared to a model reference tree. The model reference tree is a simple 4-leaf tree that best describes the relations between the 4 taxa [62, 81]. In the case of the hominid set, the reference tree has the *Pan* and *Homo* nodes as the most closely related, followed by *Gorilla* as an outgroup to *Homo*-*Pan*. For the hominin set, the Neanderthal and Denisovan are the most closely related sister groups, with *Homo sapiens* as the outgroup to the Neanderthal-Denisovan clade.

For each comparison of a generated tree with the model reference tree, we assigned a label (“Topology #1”, “Topology #2”, “Topology #3” and “Topology #4”), each corresponding to one of the four possible topologies (including a polytomy). For both sets of taxons, the four topologies and their matching labels are shown in figure 4 and figure 5. We recorded the bootstrap support value of the node with the two most closely related taxons of the generated tree, excluding polytomies. Additionally, we enumerated the number of variant sites in the concatenated alignment (excluding informative sites of the outgroup taxon) that were used to generate each tree. All protein sequences for the iterative analyses were acquired from the “Hominid Palaeoproteomic Reference Dataset”, available on Zenodo [80]. All alignments, concatenation and phylogenetic trees were generated using Module 2 of PaleoProPhyler [76]. All downstream comparisons after generating the trees were done using scripts deposited on Github (see Supplementary Material).

### Introgression

We further assessed the impact of admixture, as a contributor to apparent tree discordance. For this, we utilised the hominin dataset, given the known history of recent introgressions (genetic contributions) between the modern human, Neanderthal and Denisovan lineages [50, 64]. We first identified how often the proteins under investigation here can be found within archaic-introgressed regions of present-day human genomes. We used previously reported archaic haplotypes found within two present-day human datasets [82, 83] to assess this. The details of our methodology can be found in the Supplementary Material (S4). We also repeated the iterative analysis for the hominin and dental-bone datasets, to assess the effect of using largely un-admixed individuals when generating the phylogenetic trees. For this, we selected only individuals from the present-day human panels of the 1000 Genomes: Yoruba, Mende, Luhya and Mandinka, as the human representative. Previous studies have shown that these populations have reportedly the lowest amount of archaic introgression from the Neanderthal and Denisovan populations [82, 83].

### Phylogenetic support metrics

To better understand the relationship between the results of our analysis and the confidence metrics generated by the phylogenetic software itself, we extracted and plotted the bootstrap support of each tree from all iterative analysis datasets. We grouped the bootstrap support scores according to the data set, the number of proteins used to generate them, and the tree topology they supported. We then plotted them as boxplots using python’s 3 Matplotlib [84] package, selecting the option to not plot outliers for visual clarity.

## Results

### Incomplete lineage sorting, DNA and proteins

We inferred gene trees from each of the 12 loci of interest and observed that different topologies were recovered, depending on which data type we utilized (figure 2). For the protein set, 5 out of the 12 gene trees displayed an estimated topology that was different from the population tree (as inferred from genome-wide data [62]). In contrast, when utilizing the DNA sequences, corresponding to those 12 genes, only 2 out of the 12 gene trees differed in topology from the population tree. In all cases except two, when the protein data of a locus supported a topology different from the DNA data, the DNA data of that same locus supported the population tree topology. The two exceptions are: a) AHSG, where both DNA and protein data supported an alternative topology, which also differed from each other, and b) ENAM, where the DNA data supported an alternative topology, while the protein data supported the population tree topology.

**Figure 2:**
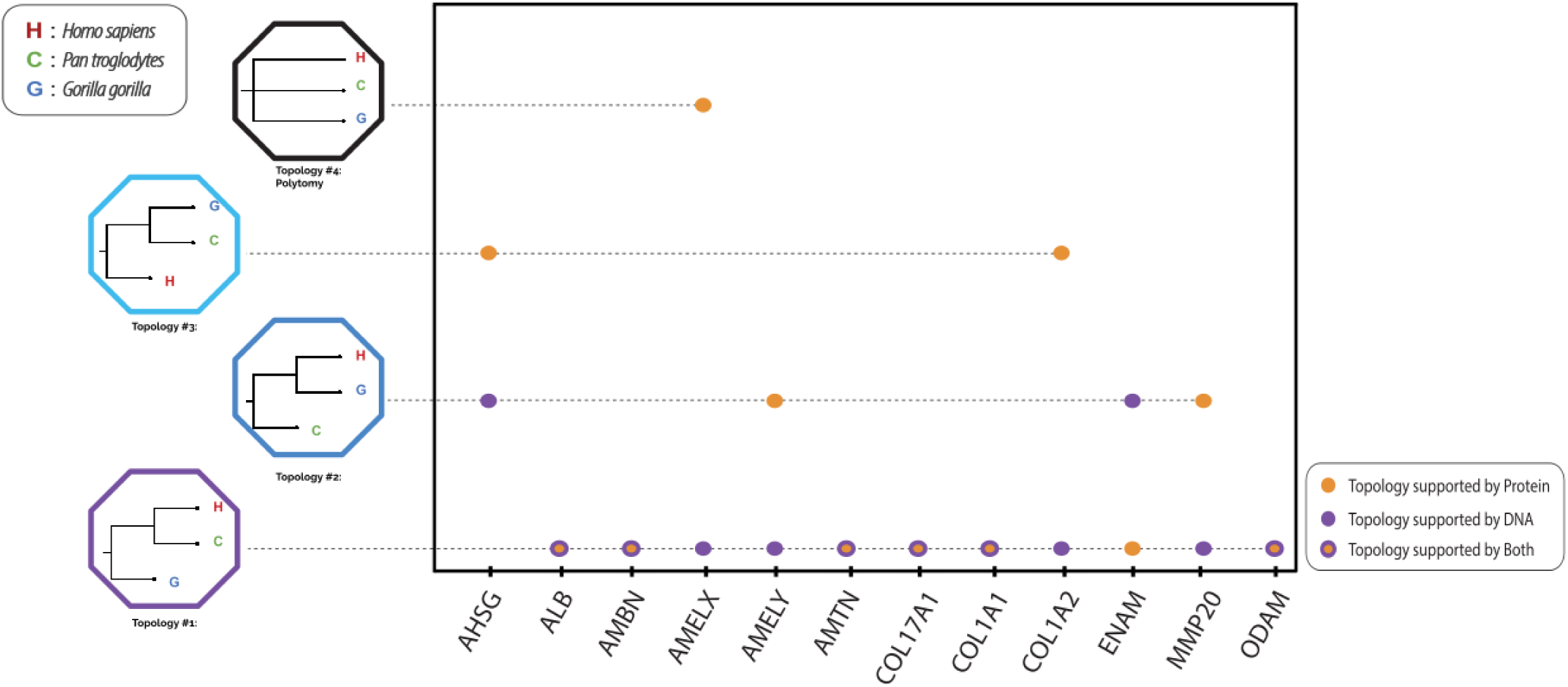
Comparison of topology supported by the 12 proteins (yellow points) under investigation and their corresponding DNA data (purple points). The 4 possible topologies are visible on the right side. For 6 loci (yellow points with purple circle), both DNA and protein data support the same topology.

### Entropy and evolutionary conservation rates

We observe notable differences between the entropy levels and evolutionary rate scores of the proteins in question (figure 3). While the aggregated entropy and evolutionary rate rankings showed slight differences, when accounting for the length of each protein, both metrics showed a nearly identical arrangement of the proteins. When not accounting for protein length, ENAM, followed by COL17A1, were found to contain the highest amount of entropy as well as the highest evolutionary rate scores. When the entropy and evolutionary rate scores were divided by the length of each protein, however, ODAM was found to be the most variable, while proteins like AMELX, COL1A1 and COL1A2, fell on the lower end of the spectrum. When compared to known conserved proteins, almost all proteins showed substantially higher entropy and evolutionary rate scores than the ubiquitin and histone proteins that we compared them with, while the fibrin proteins fall within the range of the enamel proteins.

**Figure 3:**
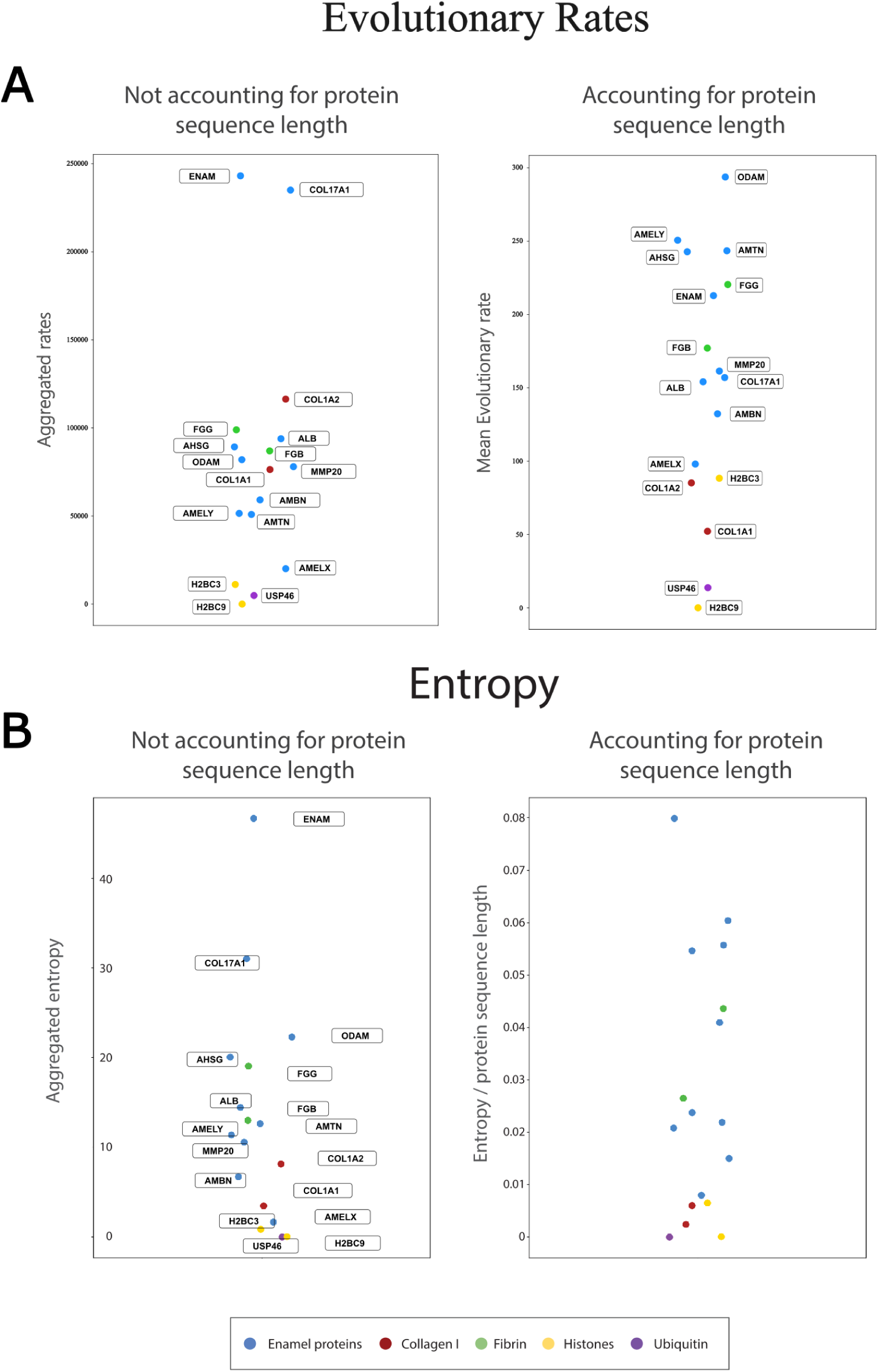
A: Protein entropy scoring comparison. Left: Each protein is ranked from highest to lowest based on the entropy scoring. Right: The entropy scoring is normalized based on the length of each protein, which causes some proteins to swap ranking. B: Protein evolutionary rates scoring comparison. Left: Each protein is ranked from highest to lowest based on the total evolutionary rate across all sites. Right: The total evolutionary rate is normalized based on the length of each protein, which causes some proteins to swap ranking.

### Informational content: exons, introns and proteins

When utilizing the entropy metric to examine the different data types, we observed that information decreased when comparing segments of mixed introns and exons to segments of pure exons and to those of amino acids (Sup. figure 1-A,B). While the difference in entropy from the mixed intron & exon data to pure exons was substantial, the difference between the exon and the amino acid dataset was much smaller in comparison. When we normalized each entropy measurement by the length of the sequence that generated it, this drop in information content did not hold. In certain cases the mixed intron & exon dataset contained less entropy than the pure exon dataset. In other cases, the amino acid alignment was the one with the highest entropy per site (Sup. figure 1-C). Differences associated with each data type’s length (triplets of nucleotides corresponding to single amino acids) may be influencing these results. To correct for this, we divided the average per-site information of the introns & exons and pure exons (seen in Sup. figure 1-C) by 3, while keeping the protein measurement intact, in order to normalise this difference in datatype length. After applying this “length-correction” to the normalised entropies, we once again observe the original pattern of informational decrease for all genes (Sup. figure 1-D), with the exception of AMELX. For AMELX, the pure exon dataset showed more information content than the mixed intron & exon dataset, even after the length-correction.

### Iterative phylogenetic analysis

Our consensus tree analysis yielded different results for the three different datasets it was applied to. For the hominid dataset, as the number of proteins and variants utilized in the concatenation increased, the number of consensus trees agreeing with the population tree topology (topology #1) increased as well, in an almost linear fashion (figure 4). At a lower number of concatenated proteins (*N* between 1-6), even if the majority of the consensus trees generated did agree with the population tree topology #1, large percentages of alternative topologies (#2, #3, #4) were also inferred. As an example, at *N* = 1, 40% percent of the consensus trees had an estimated topology that was different from the population tree (though note here that the “consensus” is just a single gene tree for *N* = 1). Topologies #2 and #3 were inferred in 20% of the iterations, with the other 20% corresponding to topology #4 (uninformative polytomy). This percentage of alternative topologies steadily decreased as *N* increased. At *N* = 9, less than 5% of the consensus topologies differed from the population tree topology. When examining each of the discordant consensus topologies, unresolved polytomies between the 3 African great apes (topology #4) were represented at lower *N* s (*N* = 1 *−* 3) but were absent at higher *N* s than 4. Above *N* = 10 almost all consensus topologies converged into Topology #1, the topology in agreement with the genome-wide DNA-inferred population tree. The number of informative sites (amino acid variants) that were used to generate each tree ranged from a mean of around 9 variants for 1 protein, to a mean of 113 variants for 12 proteins.

**Figure 4:**
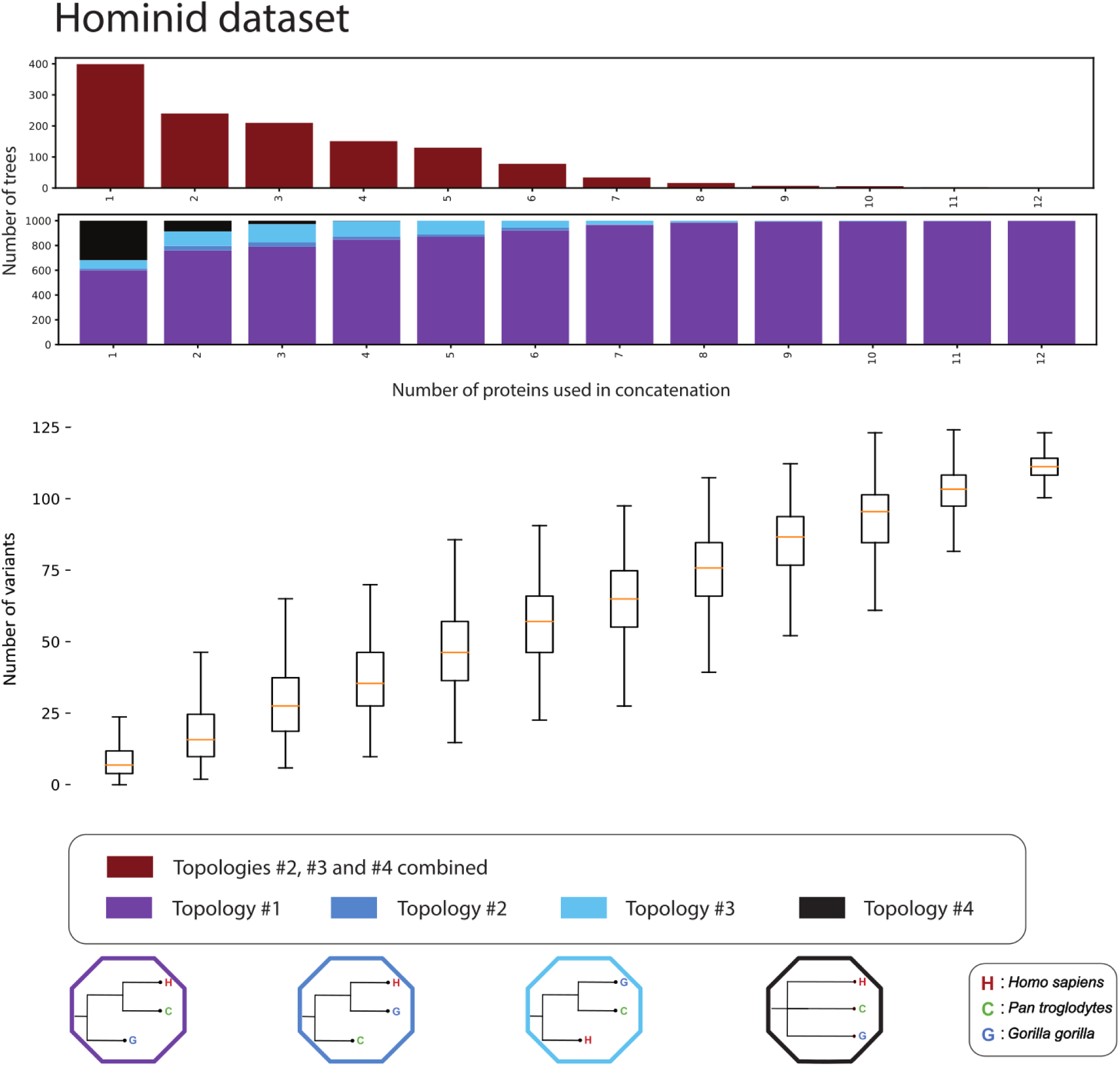
Iterative concatenation analysis for the hominid dataset. The upper barplot showcases the number of trees, out of 1000, differing from Topology #1 for an *N* number of proteins used in the concatenation. The lower barplot showcases the percentage of trees supporting each of the 4 possible topologies, out of 1000, for an *N* number of proteins used in the concatenation. The 4 possible topologies and their corresponding colours are visible at the bottom of the plot. The number of variants present in the dataset creating each tree are visible below the bar plot as a box plot. For each box the orange line denotes the median of the variants present in the *N* number of proteins concatenation. Each box plot denotes the 25%, 50% (the median, the line in the middle of the box) and 75% quantiles of the distribution. The whiskers of each box denote extremely low values (25% quantile - 1.5 * interquantile range) and extremely high values (75% quantile + 1.5 * interquantile range) for that distribution. The table containing the exact mean, median, maximum and minimum variants for each *N* proteins is available in the Supplementary Material (S3.3).

We further investigated whether the discrepancy in the amount of phylogenetic information between the enamel and collagen type I proteins can influence the generated tree topologies for the hominid dataset. To test this, we repeated the hominid tree analysis, this time excluding the 2 collagen type I proteins and using only the enamel specific ones. When doing so, we notice a slight increase in the trees in agreement with the known species tree, compared to the full enamel and collagen dataset (see Sup. figures 3 and 4). While no difference is visible when using between 1 and 5 proteins, there is a noticeable reduction in the number of discordant trees when using between 6 and 8 enamel-only proteins (or an increase in the trees supporting topology #1).

In contrast to the hominid dataset, increasing the number of proteins in the concatenation of the hominin dataset did not lead to an overall convergence to the genome-wide inferred population tree topology (topology #1) (figure 5). Instead, the number of trees supporting topology #1 remained roughly stagnant past the *N* = 5 mark, while the trees supporting topology #2 (Neanderthal and modern humans as sister lineages) steadily increased. Furthermore at *N* = 1, roughly 80% of the trees supported topology #4 (the polytomy). Although the percentage of topology #4 trees steadily dropped as the number of proteins increased, it did not completely disappear even with the use of 12 proteins (with 10% of iterations still leading to topology #4). For this dataset the number of variants was roughly 10 times lower than that of the “hominid” and ranged from a mean of 1.2 variants for 1 protein, to 14.3 for 12 proteins.

**Figure 5:**
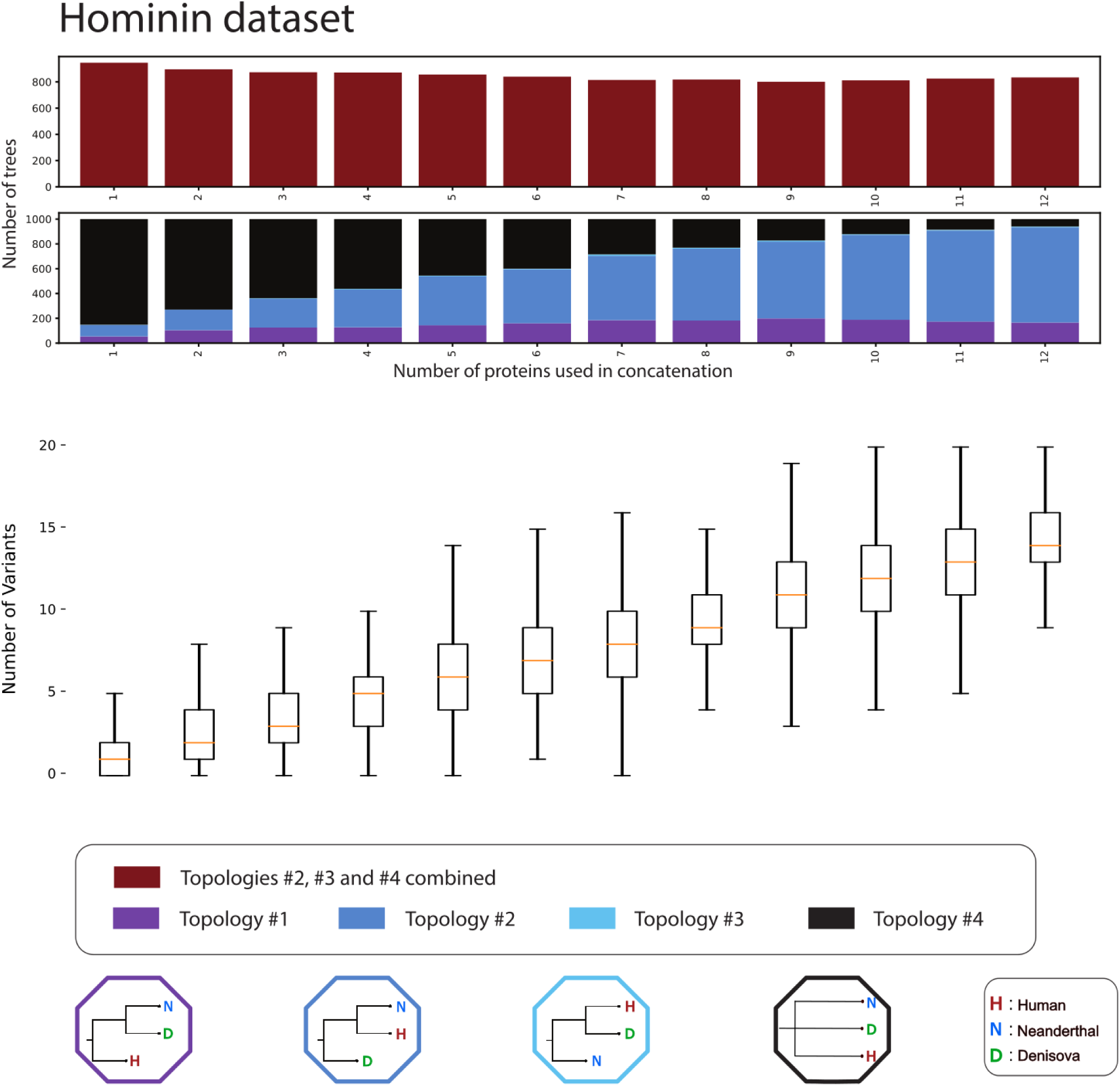
Iterative concatenation analysis for the hominin dataset. The lower barplot showcases the percentage of trees supporting each of the 4 possible topologies, out of 1000, for an *N* number of proteins used in the concatenation. The 4 possible topologies and their corresponding colours are visible at the bottom of the plot. The number of variants present in the dataset creating each tree are visible below the bar plot as a box plot. For each box the orange line denotes the median of the variants present in the *N* number of proteins concatenation. Each box plot denotes the 25%, 50% (the median, the line in the middle of the box) and 75% quantiles of the distribution. The whiskers of each box denote extremely low values (25% quantile - 1.5 * interquantile range) and extremely high values (75% quantile + 1.5 * interquantile range) for that distribution. The table containing the exact mean, median, maximum and minimum variants for each *N* proteins is available in the Supplementary Material (S3.3).

When including additional proteins (the dentin-bone dataset), we observed that the greater set of available proteins decreased the proportion of trees supporting topology #2 and increased the proportion supporting topology #1 (figure 6). In this more protein-diverse dataset, a total of 12 proteins supported either of these two topologies (#1 and #2) with roughly equal representation. Further increases in the number of proteins did not appear to change this proportion, but we did observe a continual decrease of the cases where topology #4 (polytomy) was inferred. This polytomy effectively disappeared at around 20 proteins or a mean of roughly 30 variants. At 28 proteins, or roughly 40 variants, both topology #1 and topology #2 were still equally supported.

**Figure 6:**
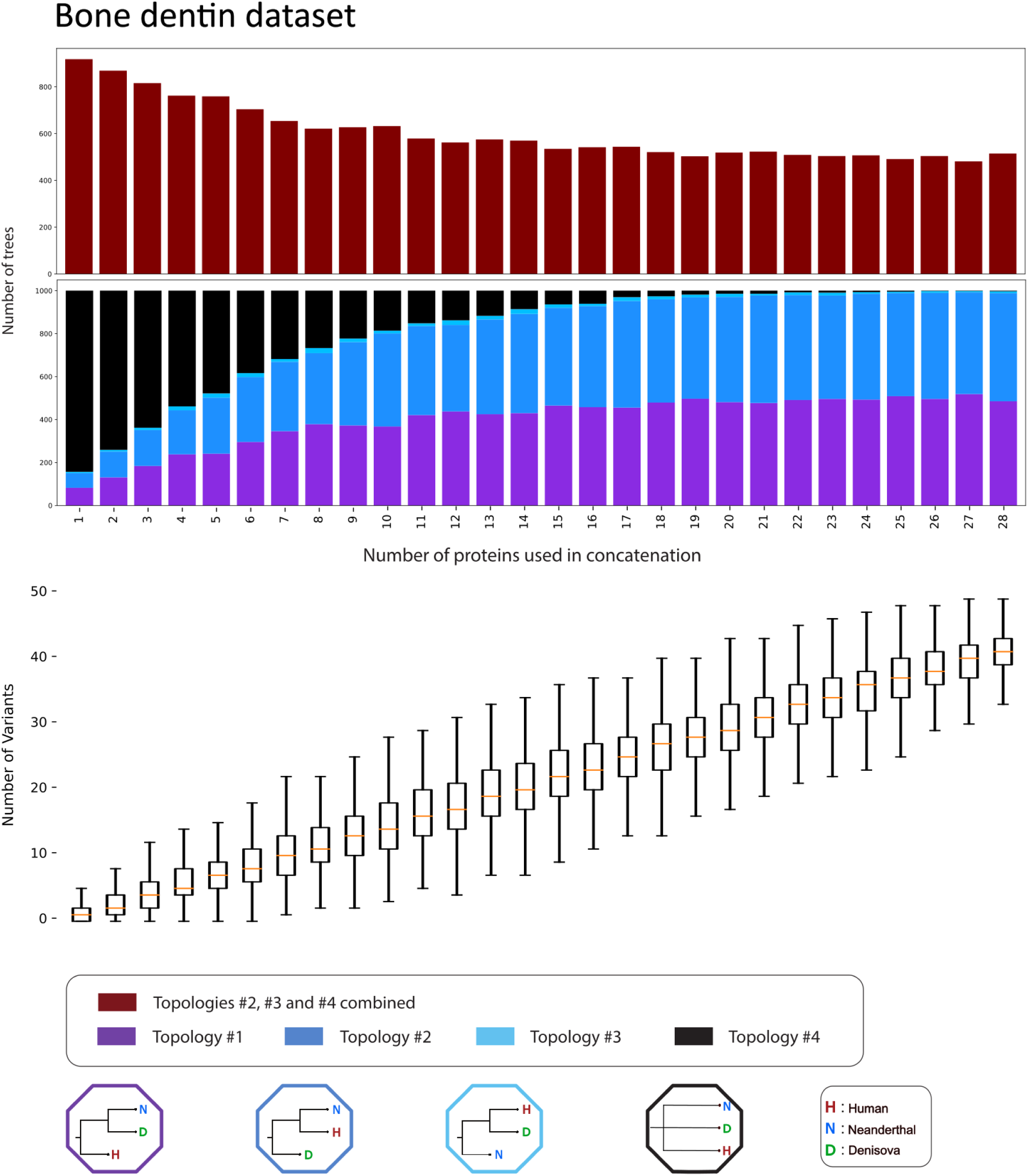
Iterative concatenation analysis for the dentin-bone dataset. The lower barplot showcases the percentage of trees supporting each of the 4 possible topologies, out of 1000, for an *N* number of proteins used in the concatenation. The 4 possible topologies and their corresponding colours are visible at the bottom of the plot. The number of variants present in the dataset creating each tree are visible below the bar plot as a box plot. For 18 each box the orange line denotes the median of the variants present in the *N* number of proteins concatenation. Each box plot denotes the 25%, 50% (the median, the line in the middle of the box) and 75% quantiles of the distribution. The whiskers of each box denote extremely low values (25% quantile - 1.5 * interquantile range) and extremely high values (75% quantile + 1.5 * interquantile range) for that distribution. The table containing the exact mean, median, maximum and minimum variants for each *N* proteins is available in Supplementary Material (S3.3).

### Introgression

Our introgression investigation showed that some of the proteins investigated here are located within archaic-introgressed regions in present-day human genomes. The frequencies of the archaic variant of a protein can be very different between each protein and between various present-day human population panels (see Sup. tables 5 - 8). In most cases, the observed frequency of the archaic-introgressed protein in the overall dataset is very low, less than 0.1%, but we found it to be very high in one of the 12 proteins: the archaic introgressed version of the enamel gene MMP20 has a frequency of around 18% in the global dataset but as high as 40% when looking at European and East Asian population panels alone. Plotting the introgressed haplotypes overlapping with MMP20 revealed a number of archaic tracks covering multiple related genes such as MMP7, MMP8 and MMP27, present in populations of almost every continent (figure 7). Other introgressed proteins showed a more localized introgression signal such as higher frequency for ALB and ODAM variants in Oceanian populations that match the Denisova 3 high coverage genome (see Sup. table 7).

**Figure 7:**
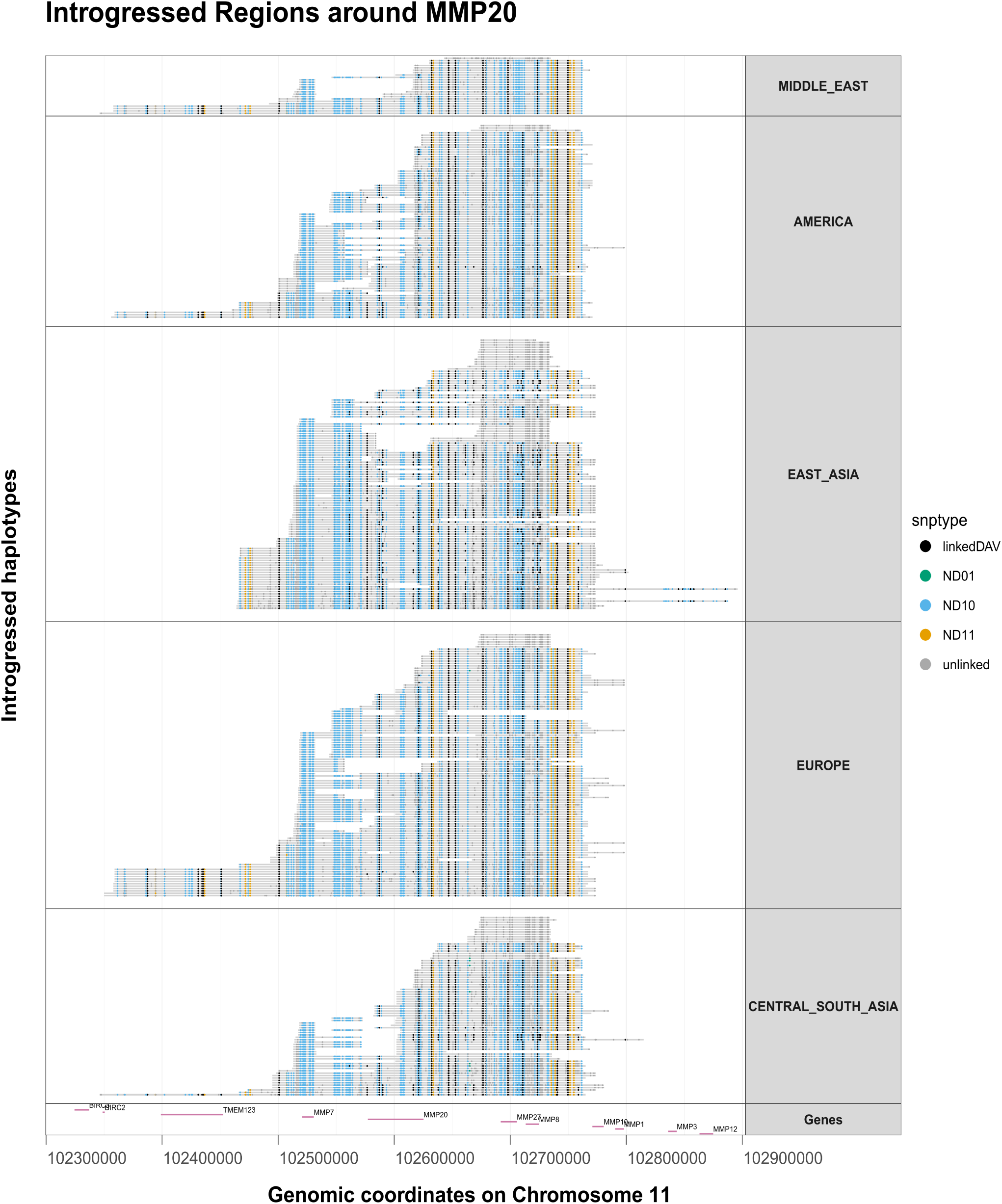
Introgressed segments (grey) overlapping the MMP20 gene and aligned to the human reference genome coordinates. Each row is an introgressed segment from a single chromosome of a specific individual. The segments have been grouped by continent and sorted by length1. 9 The dots on each segment represent single nucleotide polymorphisms (SNPs). These SNPs are either modern human derived polymorphisms that are linked (black) or unlinked (grey) to the introgressed segment, or introgressed archaic (orange), Neanderthal (blue) or Denisovan (green) polymorphisms . The last row showcases the range of the coding genes (*n* = 11) corresponding to this region, centered around MMP20.

The modified iterative analysis of the hominin and dental-bone datasets, using present-day humans only from 4 African population panels, revealed slight but noticeable differences (figure 8 and Sup. figures 11 - 13). In contrast with the original iterative analysis of the hominin dataset, when we only used African individuals from the 1000 Genomes, we observed a small but steady decline in the number of trees disagreeing with Topology #1. This included the trees generated when using the maximum of 12 enamel proteins or 28 enamel, bone and dentin proteins. Similarly, the number of trees agreeing with topology #1, while before remained stagnant past 4 proteins in the enamel dataset, now slightly increases continually until the 12 protein mark (figure 8). For the bone-dentin dataset, the results of using only African Homo sapiens had a smaller impact and once again the overall dataset supports topology #2 being as likely as topology #1. Another consequence of using only the African population panels is the complete disappearance of topology #3 (the one with humans and Denisovans closest), which may have been a consequence of removing samples from Asia harboring Denisovan ancestry (we note the 1000 Genomes Project used by our iterative analysis, has no representation of populations from Oceania, who tend to bear a considerable proportion of Denisovan ancestry).

**Figure 8:**
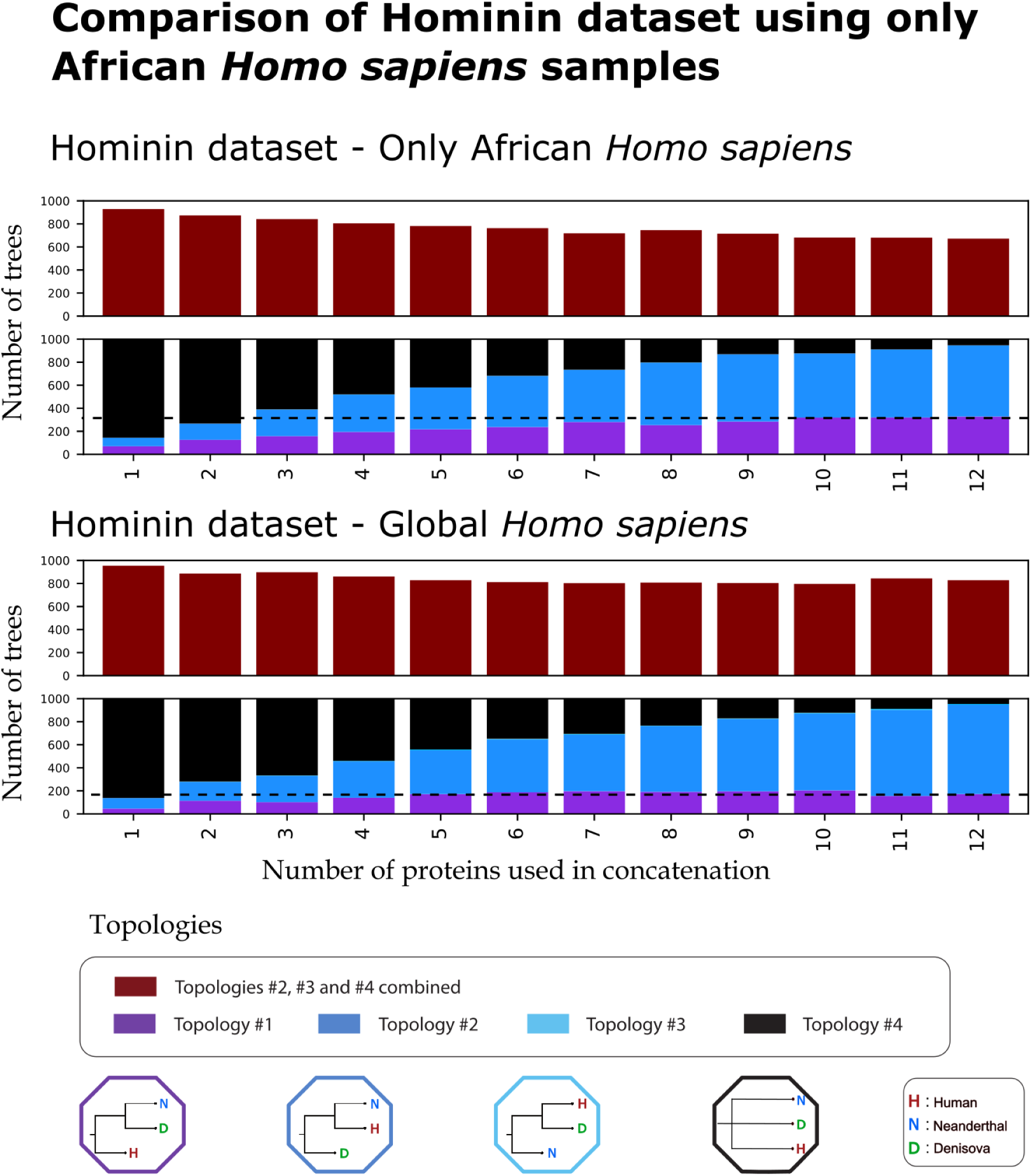
Comparison of iterative analysis results between using one of two methods for the hominin dataset a) only African samples as the Homo sapiens representative and b) a randomly chosen sample from any population of the 1000 Genomes as the Homo sapiens representative. For each of the two datasets, the first barplot at the top of the figure represents the number of trees, out of 1000 repetitions, differing from topology #1, for each number of proteins used in a concatenation, ranging from one to twelve. The second barplots breaks down the number of trees supporting each of the four topologies for the same number of proteins. A black dotted line has been added that denotes the number of trees supporting topology #1 when using the maximum number of available proteins for each of the two methods.

### Phylogenetic support metrics

Our results indicate that a high bootstrap support is a metric indicative of the robustness of the underlying protein data in supporting a topology. This is based on the fact that a mean high bootstrap support (*>*90%) was generated only in cases where the different combinations of proteins in the alignment all consistently supported the same tree, which was also matching the reference population tree. Lower bootstrap supports (*<*50%) were associated with topologies that were under represented in the total number of trees, while medium bootstrap supports (75%*<*90%) were usually indicative of multiple topologies that were presented in equal numbers. The generated figures and a detailed review of the bootstrap results are available in the Supplementary Material (S3.5).

## Discussion

Palaeoproteomic data hold immense potential for the study of hominid evolution. Ancient proteins are already being used to track the distribution of different species through space and time [22, 23, 68, 69, 85, 86, 87, 88] and to assess the taxonomic placement of specimens [10, 11, 12]. Yet so far, only a few studies have attempted to investigate the phylogenetic potential of the various recoverable ancient proteins. Buckley et al. 2014 [56] and Froment et al. 2021 [57] have previously approached this question experimentally, by sequencing bone and tooth proteomes respectively, and identifying which proteins and peptides were recoverable from the investigated tissue. They used these recovered peptides to reconstruct phylogenetic trees of different taxa, inspecting those phylogenies for accuracy (by comparing the resulting topology with known species relations), confidence (by examining the bootstrap support of the trees) and resolution (by enumerating generated polytomies). The informativeness of those proteins was also assessed, by counting the number of species-informative variants. More recently, Fong et al. 2025 [58] delved deeper into this question, using a combination of dry and wet-lab methodologies. In this work, missingness was introduced into in-silico predicted enamel protein sequences, based on experimentally observed patterns of degradation. The in-silico degraded sequence alignments were then used to generate phylogenetic trees, assessing their topological distance from a DNA-supported tree. Lastly, in a recent publication, Codlin et al. 2025 [59] translated avian egg-shell and collagen type I proteins, assessing their phylogenetic resolution and accuracy, but also highlighting the existence and possible effects of within-taxon amino acid variability.

The above studies have demonstrated some of the potential - as well as the limitations - of using palaeoproteomic data for reconstructing evolutionary relations. Questions centered around the scarcity of data, as a result of post-mortem degradation, have understandably stood at the forefront. Yet, the works described above have either focused on individual proteins or groups of proteins as a whole. In our work, we acknowledged the fact that researchers rarely have the option of choosing which proteins to utilise in their analysis, as this is usually simply the result of which peptides are recovered. As a result, we investigated the effect of incremental additions of protein data on the confidence and topology of the generated trees, exploring all possible numbers and combinations from a group of proteins. We also explored how well these protein sequences perform in reconstructing the relations of very closely related and recently admixed populations. While more challenging, these relations are often of great interest to evolutionary biology and to hominid evolution in particular. In our work we only utilized in-silico translated, complete amino acid sequences in order to simplify our analyses and comparisons with the (complete) available DNA data. We acknowledge this is never the case when working with real ancient protein data. To make the comparison more useful, we provide the number of variable amino acid positions used in each of our phylogenetic results, which can also be calculated on real data, and can be compared with the data presented here.

### Protein informational reduction and incomplete lineage sorting

Multiple studies have previously identified high levels of ILS among great apes. For example, up to 30% of gene trees in a comparison between humans, chimpanzees and gorillas have been shown to be under ILS [40, 47, 48]. Yet, the proportion of sequences under ILS drops significantly when investigating coding regions exclusively. Moreover, gene families displaying high levels of selection are even less likely to be undergoing ILS [49]. Of the 12 genes investigated here, only 2 of them (16%) showcased apparent ILS,as inferred from the DNA sequences (see figure 2). As the entropy and evolutionary rate results showcased, some of these genes are under high constraint due to selection, which may be the main driver for the relatively low level of observed ILS among them. However, when the corresponding proteins were investigated, 5 out of 12 (41%) led to the inference of a topology that was different from the population tree.

A likely explanation of this higher level of observed ILS when analyzing proteins is misestimation of the underlying gene trees, as a consequence of the reduced informational content of proteins relative to DNA. This “informational drop” between DNA and protein data types is supported by our entropy calculations, although admittedly, the question of “how much information” is contained within each data type, largely depends on how one measures said information: absolutely, per individual site or per individual codon (See sup. material S2.1-2.4). Removing the collagen type I proteins, two of the most conserved proteins of our dataset, led to a slight decrease in the number of discordant trees, when using more than 5 proteins. Similar results were also reported by [58], who eliminated specific discordant trees by removing the conserved collagen proteins from their alignments.

These results suggest that inferences based on protein data alone, may lead to higher apparent levels of ILS than what is inferred when working with DNA data. Consequentially, protein-based phylogenetic inferences may distort evolutionary conclusions and overestimate the true amount of ILS present between taxons. Previous publications on modern proteomes have made similar arguments, based on the fact that closely related species showcase ‘apparent molecular convergence’ due to inference errors from very conserved sequences [31].

### Enamel and collagen conservation

Our combined entropy and evolutionary rate results indicate that the hominid enamel proteome consist of proteins of variable levels of conservation and information content (figure 3). All protein sequences, including Collagen type I, appear more variable than the hyper-conserved ubiquitin and histone protein sequences that were used for comparison. When accounting for protein length, although some proteins like AMELX appear very conserved, all other enamel-related proteins yield greater average site variation, within hominids, than collagen type I proteins do. This pattern holds regardless of whether one measures variation through entropy or through evolutionary rate scores. As an example, ODAM displays more than 10 times the entropy score per amino acid than COL1A1 or COL1A2 and roughly three times their evolutionary rate score. These results are in line with the knowledge that collagen genes are heavily conserved in humans [89, 90]. They also agree with Codlin et al. [59], who showcased a higher conservation rate in avian collagen sequences compared to eggshell proteins, and Krueger et al. [58], who also noted the high conservation of collagens compared to enamel proteins in primates. A notable exception to this is AMELX: while being the most abundant enamel protein [91], it shows a similarly conserved sequence to collagen type I. Yet, while AMELX tends to display low sequence variation, its Y chromosome isoform, AMELY, is much more variable, especially when accounting for its short length. This should not be unexpected: AMELY is located on the non-recombining region of the Y chromosome where it is evolving under less selective constraint and under a faster local mutation rate [92, 93, 94].

Our work is an initial investigation into the informativeness of AMELY, a protein that has recently become of great interest [95, 96, 97, 98, 99, 100, 101] due to its ability to identify the biological sex of heavily degraded samples [102, 103, 104]. Although it is expressed at lower concentration than AMELX [105], AMELY can provide useful information for species identification and phylogenetic inferences, due to its high variability. However, analyses based on AMELY are also fraught with difficulties. For most vertebrate species, even when the sequence of AMELX is well characterized, the amino acid sequence of AMELY and the location of its coding gene are unknown [70, 106]. In some taxonomic groups, the gene responsible for expressing AMELY is missing in its entirety [107]. Finally as noted by Fong et al. 2025 [58] (who for practical reasons chose not include AMELY in their enamel protein investigation), in some taxons the genes of AMELX and AMELY are not acting as independent loci [108, 109], limiting their phylogenetic utility. As a result, while our anaysis here showcases the high informativeness of AMELY - in contrast with that of AMELX - we recommend caution when working with this protein for evolutionary inferences.

Today, protein-based archeological species identification primarily relies on collagen (type I) mass fingerprinting [5, 110]. Our results indicate that, as a whole, the enamel proteome evolves faster than collagen type I and thus, when the appropriate tooth tissue is available, could differentiate between more closely related populations or species. This is especially important given the micro-destructive techniques, such as acid etching, that have successfully been applied to tooth enamel and bone material [111, 112, 113, 114]. These methods can extract useful amino acid sequence information using a minimal amount of material, inflicting only minor surface damage but preserving morphological information.

### Number of proteins and phylogenetic resolution

The 12 proteins investigated here have previously been used in different combinations and have been shown to discriminate between the 4 extant genera of the hominidae family [10, 11, 13]. Nevertheless the exact number of proteins required to reliably infer relations between these species is not yet clear. Our iterative analysis on the hominid dataset showed that, as one might expect, the number of consensus trees supporting an alternative topology to that of the population tree, drops significantly with the inclusion of additional proteins in the analysis. An increasing number of proteins sees an overall drop in discordant trees, in a linear decay, up to around 9 or 10 proteins, which we conclude to be sufficient for consistently recovering the reference population tree. Additionally, a combination of any two of the twelve proteins drops the percentage of polytomies from the population tree from 30% (when using only a single protein) down to less than 20%, and a combination of four protein, to less than 1%. Thus, simply distinguishing between these 3 groups (without accurately inferring their phylogenetic relationships), should be possible with the recovery of around 4 of any of these proteins. We expect other taxa with similar genetic distances, as those between the extant genera of hominidae, to have similar phylogenetic resolution and power using palaeoproteomic data.

For our hominin dataset instead, our analysis revealed that the number of consensus trees agreeing with the topology of the population tree, did not increase with the inclusion of additional protein sequences in the analysis. Steadily increasing the number of these proteins from 4 to 12 showed little to no improvement in resolving this phylogeny. Instead, increasing the number of proteins from this set led to an increase in support for one of the 3 alternative topologies, topology #2, the one with modern humans and Neanderthals as sister lineages. Indeed, this one particular topology (itself discordant relative to the population tree) is supported by the fully concatenated 12-protein dataset we chose for this analysis (also shown previously by Welker et al. 2020 [10]).

The inclusion of additional proteins from the bone-dentin dataset led to slightly different results. Instead of a single topology that doesn’t match the population tree becoming increasingly supported, two competing topologies (topology #1 and topology #2), equally represented, become the most commonly observed when using between 10 and 28 proteins. Similarly, a polytomy between these three groups was still present when using up to 20-24 proteins, although admittedly in extremely low frequency. Our analysis reveals that when examining very closely related populations with limited protein data, the addition of a few more protein sequences may not always increase support for the tree that is closest to the population tree of said taxons, as inferred from genome-wide data. In such cases, trees with medium to high bootstrap values (70-80%) may be obscuring incongruence in the underlying data. Likewise, distinguishing between these groups, a process necessary for species or population identification, is also a difficult task, largely dependent on recovering some of the few informative sites that exist.

One explanation for the above results is that the recent admixture between these groups [50, 64], has led to some present-day humans carrying Neanderthal or Denisovan haplotypes that overlap with the genes coding the proteins under investigation. Here we showed that a) some present-day humans do carry the archaic-introgressed version of the studied proteins, in frequencies which also differ among populations, b) that controlling for this introgression by using unadmixed present-day humans in the phylogeny does increase the proportions of the trees that are in agreement with the whole genome data and c) even when controlling for archaic admixture, the concatenated protein phylogenetic trees result in different alternative topologies with equal support for these 3 groups.

Other forms of admixture could also be influencing these results. Previous publications have hypothesised about a deeply archaic introgression into Denisovans [66], which would make this population more different than the present-day human or Neanderthal lineage. Alternatively an earlier introgression of ancient African, anatomically-modern humans into Neanderthals [83, 115], would also bring these two groups closer to each other than to Denisovans. Both such admixtures could help explain why the protein data support the topology of present-day humans being closer to Neanderthals (see figure 5 and figure 6).

Another explanation to this issue is that the fairly recent split between these 3 groups, estimated by some to be around 400.000 to 600.000 years ago [66], does not allow for accurate phylogenetic inference using the phylogenetically-conserved protein data. Given the slow evolutionary rate of protein sequences in general, it is possible that not enough time has passed for these sequences to sufficiently differentiate from one another. To put things in perspective, when comparing the number of variants present in the alignments of all 12 enamel proteins of hominids to that of hominins, the former is roughly 10 times higher than the later.

### Closing remarks and future prospects

Currently, enamel and collagen type I proteins remain the only phylogenetically informative biomolecules that are recoverable for fossil taxa in deep time (samples that are more than 1 million years old). Although this unique resource is unparalleled in terms of preservation [8, 24], its phylogenetic potential may be more limited than previously thought [10]. In homininds, many studies have already noted a lack of resolution at finer taxonomic levels: unresolved polytomies generated using the enamel proteome have been identified inside the genus of *Pongo* (between the 3 extant species) [11, 12], the genus *Gorilla* (between the 2 extant species), as well as within the genus *Homo* (between present-day humans, Neanderthals and Denisovans) [6, 10, 88]. Polytomies have also been observed at a subspecies level, such as the divisions between subspecies of *Gorilla gorilla* and of *Pan troglodytes*.

Nevertheless, the two species of the genus *Pan* (*P. troglodytes* and *P. paniscus*) can be confidently distinguished from one another using a concatenation of enamel proteins [10, 11]. The reason why some evolutionary relationships are easier to resolve than others needs to be further investigated, but probable causes include differences in split times, differences in the effective population size of ancestral populations and different levels of post-divergence migration [43].

Both results from previous publications and the present study suggest that phylogenetic analyses of archaic hominid taxa based on palaeoproteomic data should be taken with a degree of caution. Overall, protein data may lead to higher amounts of gene tree misestimation, as a result of the data type used for tree estimation. Here we have shown that the number of currently recoverable, deep-time proteins allows for the reconstruction of species relations at the level of genera in the hominid clade. This is very encouraging, given this particular clade’s genetic history of recent splits, high levels of ILS [40, 47, 48, 49] and past admixture events [50, 51, 52, 53]. However, our results also indicate that the same data have limited power to resolve the population trees of more closely related groups, such as those within the hominin clade.

The issues described here are neither new nor unique to the field of palaeoproteomics. During the early decades of the field of molecular phylogenetics, the limited amount of sequence data at the time, initially proteins, and later on short DNA sequences, offered limited resolution when resolving clades of closely related species. As an example, early studies were unable to resolve the polytomy of the human, chimpanzee and gorilla lineages [116] and identify which species was our closest living relative. This issue was not resolved until the accumulation of sufficient data roughly two decades later [117]. Similarly, early ancient DNA studies based only on mitochondrial DNA, supported a scenario of “no admixture” between Neanderthals and modern humans [118, 119]. The first published mitochondrial DNA from a Denisovan, characterized them as an outgroup to Neanderthals and modern humans [120]. Once again, these relationships were reconsidered and resolved with the acquisition of more molecular data. Increases in overall ancient peptide acquisition through novel lab methodologies [68, 121, 122, 123] may lead to more confident phylogenetic placements and enhanced evolutionary resolution for these taxa. Studies extracting the bone proteome of younger samples have so far delivered a greater number of proteins [15, 16, 18, 19, 21]. As an example, two recent publications have managed to recover an impressive amount of bone and dental proteins (*n* = 51 [22], *n* = 88 [23]), which they used to phylogenetically assign two fossil specimens to the Denisova clade, expanding our understanding of this enigmatic group. Future advances, such as targeted proteomic approaches [100, 124], the identification of better preserving bones [125] or improvements in downstream spectra identification [126, 127] may allow for similar recoveries in older samples. However, we believe that an even greater number of proteins than the ones investigated here (*n* = 28) will be necessary for the accurate resolution of evolutionary relations for very closely related populations or species. Alternatively, more computationally-intensive methods of inference beyond concatenation, such as the multispecies coalescent (MSC), could result in better resolution by better accounting for the evolutionary processees that lead to different gene trees along a sequence [128, 129]. Given that the protein data examined here features high incongruence and a low number of informative loci, it is possible that tools like *Beast [130] might provide results that better agree with the evolutionary relationships inferred from DNA.

On top of this, the population tree itself may be a poor representation of the overall relationships between closely related groups, due to admixture events [50, 64, 65, 66]. Indeed, when investigating the relationships between organisms that are as closely related as the ones investigated here, concepts such as “species” or “trees” lose some of their utility. A growing body of work from the field of ancient population genetics has shown that admixture between even distantly related groups of hominids might be the standard rather than the exception [51, 52, 131]. This seems to be especially true within the confines of hominin evolution during the Late Pleistocene [4, 132, 133, 134]. In light of these discoveries, the field of paleoanthropology is also changing. Past quests for a single population tree are now slowly being replaced by the concept of a “braided stream”, a network of reticulating lineages that can split as much as they can merge [3, 134, 135]. Reconstruction of hominin evolutionary relations using other topological objects beyond trees (like admixture graphs) is still in its infancy, and so far, non-existent in the field of paleoproteomics. This, in turn, suggests a fruitful avenue for future methodological developments.

## Data Availability

The data and scripts to reproduce the entropy calculations between the different proteins and other data types (exons-only, exons-and-introns), along with the script to reproduce the results of figure 2, are available on Zenodo: https://zenodo.org/records/17530636 [136]

The scripts and data, and download links to reproduce the iterative tree analysis, as well as the introgression investigation are available on Github: https://github.com/johnpatramanis/Protein_ILS_Hominids_and_Hominins [137].

## Acknowledgements

We thank Alberto John Taurozzi, Evan Irving-Pease, Graham Gower, Martin Petr, Johanna Krueger, Ryan Sinclair Paterson and other members of the Racimo and Cappellini groups, who provided valuable help, suggestions and feedback throughout the project. We also want to thank Professor Alan Rogers, Prof. Fernando Villanea and the anonymous referee for openly reviewing our manuscript and providing useful recommendations, ideas and suggestions.

