## Supplementary Material for "Assessing the potential of ancient protein sequences in the study of hominid evolution"

### Supplementary Methods and Results

#### 1. DNA and protein apparent incomplete lineage sorting

##### 1.1 The Dataset

The reference sequences for the following 12 proteins were acquired from Ensembl [1]: AHSG, ALB, AMBN, AMELY, AMELX, AMTN, COL17A1, COL1A1, COL1A2, ENAM, MMP20 and ODAM and for the following four hominid species: *Homo sapiens*, *Pan troglodytes*, *Gorilla gorilla*, *Pongo abelii.* The Ensembl ID's for these sequences are available below (Table 1). The ortholog sequences from the four hominid species were aligned using Mafft [2]. Gene trees were reconstructed using PhyML [3] for each of the 12 ortholog alignments separately. The 12 generated trees were rooted using *Pongo abelii* as the outgroup and compared to the population tree that best represents the relationships between those 4 species [4]. Our protein tree results were compared to those using genetic data, by repeating the same process using the reference DNA sequences (both exons and introns) of the genes corresponding to those 12 proteins. Both DNA level data and protein data are available at https://zenodo.org/records/17530636 [5].

| | |
|---|---|
| *Homo sapiens* | ENST00000411641.7, ENST00000295897.9, ENST00000322937.10, ENST00000651267.2, ENST00000380712.7, ENST00000339336.9, ENST00000225964.10, ENST00000648076.2, ENST00000396073.4, ENST00000260228.3, ENST00000683306.1,ENST00000297268.11 |
| *Pan troglodytes* | ENSPTRT00000029292.6, ENSPTRT00000064657.3,ENSPTRT00000045270.2, ENSPTRT00000065610.3, ENSPTRT00000084033.1, ENSPTRT00000030049.4, ENSPTRT00000017231.5, ENSPTRT00000005563.4, ENSPTRT00000091821.1, ENSPTRT00000007863.3, ENSPTRT00000030041.4, ENSPTRT00000035921.4 |
| *Gorilla gorilla* | ENSGGOT00000047574.1 , ENSGGOT00000024652.2 , ENSG-GOT00000013709.3, ENSGGOT00000023433.2, ENSGGOT00000016458.3, ENSGGOT00000013270.3, ENSGGOT00000000517.3, ENSG-GOT00000017106.3, ENSGGOT00000003330.3, ENSGGOT00000007947.3, ENSGGOT00000006068.3 |
| *Pongo abelii* | ENSPPYT00000016730.3, ENSPPYT00000055783.1, ENSPPYT00000017210.2, ENSPPYT00000023465.2, ENSPPYT00000017209.2, ENSPPYT00000010431.3, ENSPPYT00000003164.2, ENSPPYT00000017211.2, ENSPPYT00000004535.2, ENSPPYT00000017201.2 |

Table 1: Table matching each of the four species with the Ensembl reference ID of the 12 genes under investigation.

##### 1.2 Reproducing the results

###### 1.2.1 Setup

All the necessary code and data to reproduce this analysis are available at Github [6]:

https://github.com/johnpatramanis/Protein_ILS_Hominids_and_Hominins/tree/main

and require only a Linux machine with conda installed to run. To reproduce the code however, the user should first install the 2 conda environments that make all the tools available. The two environments are available in the main folder and can be installed with the following commands:

```
#### Install "Analyser" environment
conda env create -f Analyser.yml

#### Install "Entropy" environment
conda env create -f Entropy.yml
```

##### 1.2.2 DNA Enamel Hominid Gene Trees

To rerun the DNA trees for the 12 enamel genes, simply download the repository from Github:

https://github.com/johnpatramanis/Protein_ILS_Hominids_and_Hominins/tree/main

and enter the "ILS - Hominids/Enamel_Hominids_Trees/Enamel_DNA_Hominds_Tree_Distances" folder with your command line. Then type:

```
### Activate environment
conda activate Analyser

### Move into the relevant directory
cd GENE_TREES_ENAMEL

### Execute Script
bash Generate_Trees_PhyML.sh

### Deactivate Environment
conda deactivate

### Return to main folder
cd ..
```

This will generate a single tree for each gene and format the label names for you. To then compare each of these 12 trees with the reference topology, simply execute:

```
### Activate environment
conda activate Entropy

### Execute R Script
R Tree_Dist.r

### Deactivate Environment
conda deactivate
```

This should generate a "Hominid_GENENAME_Tree_Distances.txt" distance file, inside the GENE_TREES_ENAMEL folder, for each gene. The first metric of the distance file denotes the topology supported, e.g. “2” denotes topology number 2, or “T2”. The second number denotes the pure distance from the reference tree, with for example “1” being a different topology and “0” being an identical topology. The remaining numbers in the distance file denote alternative distance metrics generated, but where not utilised in any way.

##### 1.2.3 Protein Enamel Hominid Gene Trees

To rerun the Protein trees for the 12 enamel genes, download the repository from Github:

`https://github.com/johnpatramanis/Protein_ILS_Hominids_and_Hominins/tree/main`

and enter the folder "`ILS - Hominids/Enamel_Hominids_Trees/Enamel_Protein_Hominds_Tree_Distances`" with your command line. Then type:

```
### Activate first environment
conda activate Analyser

### Execute Script
bash Generate_Trees_PhyML.sh

### Deactivate Environment
conda deactivate
```

This will generate a single tree for each protein. To then compare these 12 generated trees with the reference topology, simply execute:

```
### Activate second environment
conda activate Entropy

### Execute R Script
R Tree_Dist.r

### Deactivate Environment
conda deactivate
```

Just like in the previous example (1.2.1), this should generate a "`GENENAME_Tree_Distances.txt`" distance file inside each GENE folder. The first metric of the distance file denotes the topology supported, e.g. "2" denotes topology number 2, or "T2". The second number denotes the pure distance from the reference tree, with for example "1" being a different topology and "0" being an identical topology. The remaining numbers in the distance file denote alternative distance metrics generated, but where not utilised in any way.

##### 1.3 Plotting the results

After running the code provided in 1.2.1 and 1.2.2, you can now also generate the main component of Main text figure 1 by entering the `"Enamel_Protein_Hominds_Tree_Distances"` folder and executing:

```
### Run Python Plotting script
python3 Topology_Per_Gene.py
```

Note that there is no environment provided for the plots and executing the above script requires that you have python3 installed with the package of Matplotlib.

#### 2. Entropy and Evolutionary Rates Calculations

##### 2.1 Entropy Comparisons between data types

In order to assess the phylogenetic information in the proteins studied here, we utilised a metric of “entropy”. This metric is based on Shannon’s information theoretic entropy measurement [7], as implemented by the Bio3d R package [8]. To assess the efficacy of this measurement in calculating phylogenetic information, we quantified the informational loss between the three data types of a) combined introns and exons, b) pure exons and c) amino acids. We assembled 12 alignments, one for each gene under study, for each of the 3 data types. Each alignment contains the ortholog sequences for 4 hominid species: *Homo sapiens*, *Pan troglodytes*, *Gorilla gorilla* and *Pongo abelii*. The sequences were obtained by randomly selecting one individual for each species from the Great Ape Genomes project [9]. All data types (combined exon and intron data, pure exons and translated amino acid data) were generated directly from the BAM files [10] using PaleoProPhyler’s translation module [11]. The original bam files were generated by mapping the raw fastq files onto the GRCh38 human reference genome and are available upon request. The ortholog sequences were aligned using Mafft [2]. The 36 ortholog alignments (12 genes x 3 data types), organised in 3 datatype specific data sets are available at `https://zenodo.org/records/17530636`[5]. We applied Bio3d’s entropy metric to each of the 36 ortholog alignments, acquiring an entropy measurement for each position of the alignment. We then aggregated the per position entropy into a single measurement and compared it across the different genes and the different data types (Main text figure 3). A version of the same plot, but with better resolution on all values lower than 4000 is available below (Sup. figure 1). We then also normalised that number by dividing with the total length of each data type. This new measurement (Main text figure 4) represents the average information per site of each data type (per nucleotide for DNA data, per amino acid for protein data).

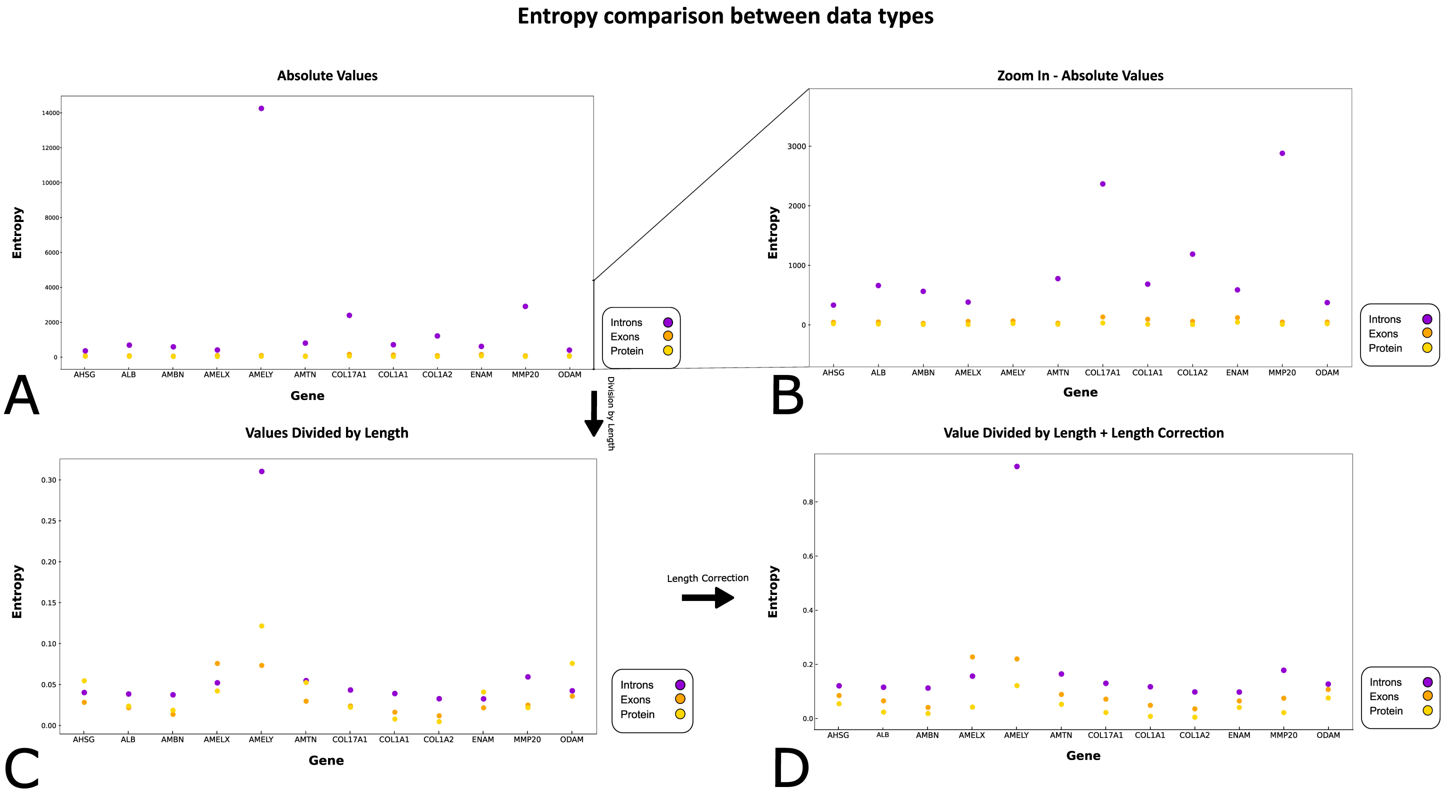


Figure 1: Entropy scoring comparison between each locus in three data types: Introns (mixed introns and exons), Exons and Protein. A: The absolute entropy values from each position of the data type (either nucelotide or amino acid site) are summed together into a single value, which is plotted without any alterations. B: Zoom-In of absolute values to better distinguish data types that have very similar absolute values. C: The summed absolute value is divided by the total length of the datatype (e.g. number of bases). D: The summed absolute value is divided by the length of the datatype, but then the protein entropy is further divided by 3, to bring it into line with DNA datatype.

##### 2.2 Reproducing the results - Entropy Comparisons between data types

All the necessary code and data to reproduce this analysis is available at Github:

`https://github.com/johnpatramanis/Protein_ILS_Hominids_and_Hominins/tree/main`

and requires only a Linux machine with conda installed. To reproduce the code however, the user should first install 2 conda environments that make all the tools available. The two environments are avaialble in the main folder and can be installed with the following commands:

```
### Install environment
conda env create -f Entropy.yml
```

###### 2.2.1 Data Type Entropy Analysis

To reproduce the data type entropy analysis, first download the repository from Github:

`https://github.com/johnpatramanis/Protein_ILS_Hominids_and_Hominins/tree/main`.

Then navigate using the command line to the :

`Entropy/Comparison_Exons_Introns_Proteins/Homind_Data_Type_Entropy_ALIGNMENTS/`

folder. There, simply execute:

```
#### Activate Environment
conda activate Entropy

### Execute Script
bash Calc_Entropy_Introns_Exons_Proteins.sh

### Deactivate Environment
conda deactivate
```

This will generate a single file ending with ".entr", containing the entropy metrics, for each of the 36 alignments and place those files into the `"Entropy_Measurements"` folder.

##### 2.2.2 Plotting the results of Data Type Entropy Analysis

After generating the entropy files from 2.2.1, you can also generate the plots (Main text figures 3,4) simply by moving up a folder into `Entropy/Comparison_Exons_Introns_Proteins/` and executing:

```
### Run Python plotting script
python3 Plot_Entropies_IEP.py
```

Note that there is no environment provided for the plots and executing the above script requires that you have python3 installed with the package of Matplotlib.

##### 2.3 Entropy and Evolutionary Rates Calculations per protein

To more accurately compare the amount of phylogenetic information between these 12 proteins we designed an analysis that also takes into account the within species protein diversity. For this analysis, we utilised the Hominid Palaeoproteomic Reference Dataset (HPRD), available from https://zenodo.org/records/7728060, which provided us with a range of individuals for each species for all 12 proteins analysed. Our analysis uses an iterative model. For each of the 12 proteins, we first randomly select one individual from each species from the dataset and isolate from them the protein of interest. We used no aligner since the protein sequences from the four different species are already aligned in the HPRD. We then apply the same entropy calculation that is described in 2.1 to the 4 species alignment and store the 2 entropy metrics generated (mean entropy per site of the protein and flat entropy across the whole protein). Additionally, we apply Rate4Site's [12] evolutionary rate calculation, by running "rate4site -s Input-Alignment-File -o Output-File". We then repeat this process for 1000 repetitions, each time re sampling with replacement four individuals from the dataset. We then calculate the mean from all repetitions using the stored measurements and end up with two metrics of entropy and two metrics for the evolutionary rate, for each of the 12 proteins. To contextualise our results we also added 5 proteins that have been previously reported as being either very conserved or containing hyper-variable segments. The same calculations is applied to these alignments as well, but only for a single repetition, using the reference data from Ensembl (these proteins were not available in the HPRD). These 5 proteins included two histones (H2BC3,H2BC9) and one ubiquitine (USP46) as "conserved" proteins, as well as two "variable" fibrinogen proteins (FGB,FGG).

**A note on long tree branches:** In some particular cases of multi-species alignments, its possible that one of the species has dis-proportionally more mutation than the other species. An example of this is AMELX, where humans, chimpanzees and gorillas have identical sequences, but orangutans differ from them at two sites. When generating a tree from this kind of alignment, one of the branches of the tree (in this case the orangutan branch), will showcase a much longer branch. When calculating the entropy for these alignments, one will also get a measurement that implies diversity within the alignment, however this diversity comes only from one (or some) of the branches. In order to try to correct for that we devised a modified entropy metric, which we will be calling 'long branch correction entropy'. Instead of calculating a single entropy for a four species alignment, we calculated the entropy of all unique pairs of the alignment and then averaged that into a single entropy. Similarly to before we calculated both a flat corrected entropy as well as an average entropy divided by the length of the alignment. We incorporated this new entropy metric and calculated it at the same time as with the original entropy. Although these corrected entropies are consistently lower than the original ones, the order of proteins does not change at all figure 2. Due to this, we did not incorporate this entropy metric into the main text, nor did we try it on the evolutionary rates.

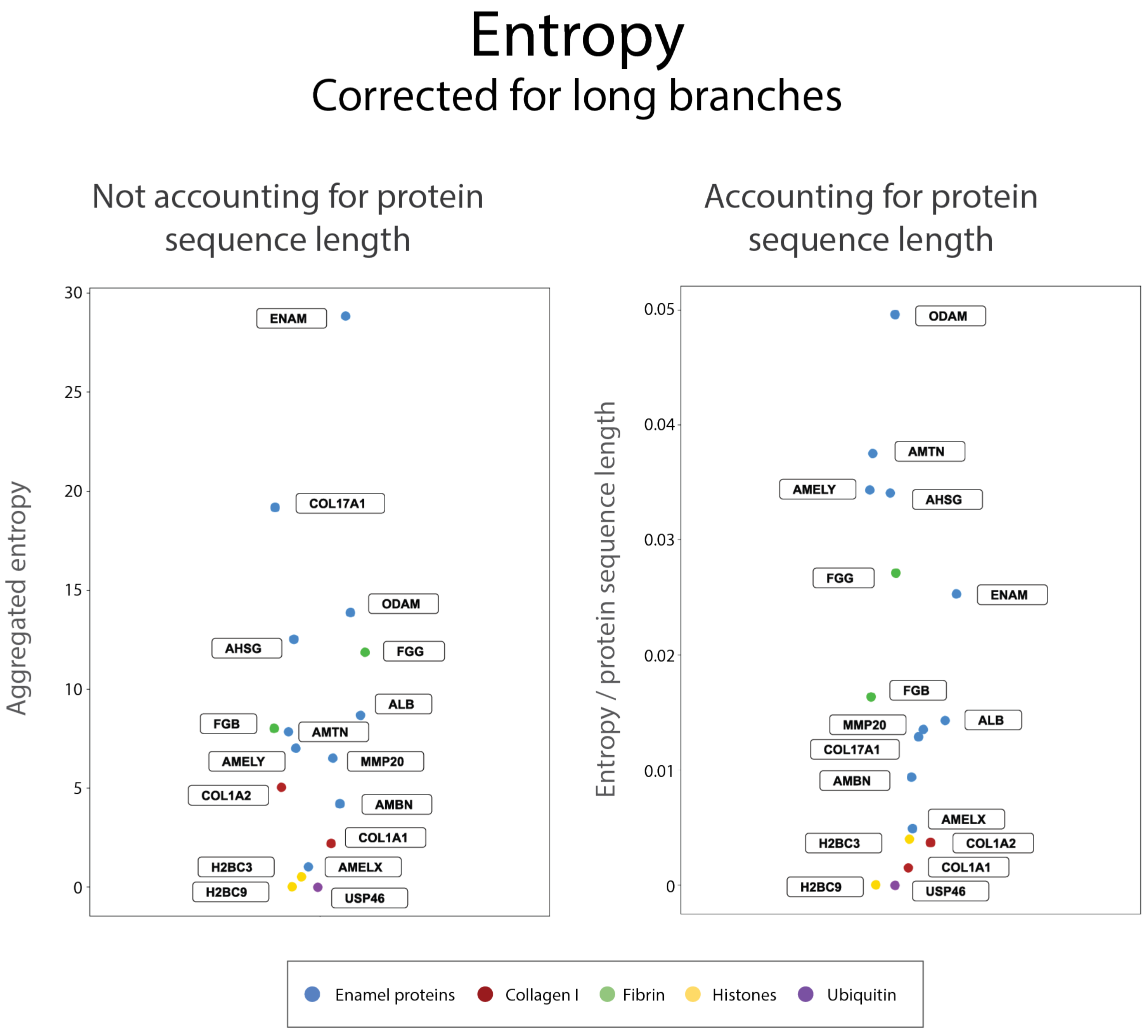


Figure 2: Protein entropy scoring comparison, with correction for longer branches on the tree. Left: Each protein is ranked from highest to lowest based on the entropy scoring. Right: The entropy scoring is normalized based on the length of each protein, which causes some proteins to swap ranking.

##### 2.4 Reproducing the results - Entropy and Evolutionary Rate Calculations per protein

All the necessary code and data to reproduce this analysis is available at Github:

`https://github.com/johnpatramanis/Protein_ILS_Hominids_and_Hominins/tree/main`

and require only a Linux machine with conda installed. To reproduce the code however, the user should first install 2 conda environments that make all the tools available. The two environments are available in the main folder and can be installed with the following commands:

```
#### Install Environment
conda env create -f Entropy.yml
```

###### 2.4.1 Entropy and Evolutionary Rate Calculations per protein

To rerun the Data Type Entropy Analysis described in the previous section, all you need to do is download the repostiory from here: `https://github.com/johnpatramanis/Protein_ILS_Hominids_and_Hominins/tree/main` move using your command line into the `Entropy/` folder and then simply execute:

```
##### Activate Environment
conda activate Entropy

#### Run workflow of multiple scripts
snakemake -j4

##### Deactivate Environment
conda deactivate
```

Note that instead of using snakemake -j4, one can use any number of cores their computer provides them with. E.g. a computer with 32 cores can run snakemake -j32. This will make the analysis run faster, as it is heavily parallelised.

The above command will generate a single file per gene/protein analysed named `"GENE.entr"`, where GENE is the name of each protein. These folder will be placed inside the `'Entropies_Hominids'` folder, along with a file named `"Combined_Entropies"`. The individual GENE.entr files should contain the resulting entropies from 1000 iterations. The numbers in each row (each iteration), correspond to: a) The mean entropy per amino acid of the protein, b) The flat entropy of the protein, c) The flat entropy of the protein, corrected for longer branches, d) The mean entropy per amino acid of the protein, corrected for longer branches. For a more detailed explanation on each of these entropies, please read above. Additionally this folder will contain one subfolder for each GENE. Within each of these subfolder are contained each 1000 output files (IterationNumber.output) for each iteration of Rate4sites run.

The average entropy (mean, flat, mean corrected, flat corrected) for each protein after 1000 repetitions is written out in the `"Combined_Entropies"` file, while the average evolutionary rate (flat, mean) after 1000 repetition is written out in the `"Combined_Rates"` file.

### 3. Iterative Analysis

#### 3.1 Effect of number of proteins in tree inference

##### 3.1.1 Detailed methodology of iterative analysis

To investigate the effect of using of different numbers and combinations of the available proteins on the generated phylogenies, we designed an iterative analysis. We applied this analysis on to two different quadruples of organisms. The first set, mentioned here as “hominid dataset” consisted of the species: *Homo sapiens*, *Pan troglodytes*, *Gorilla gorilla* and *Pongo abelii.* The second set, mentioned here as “hominin dataset” consisted of the following 4 groups: *Homo sapiens*, Neanderthals, Denisovan and *Pan troglodytes.* For each set, one individual from each group was selected randomly from the HPRD to represent the group. An $N$ number of proteins, sampling randomly and without replacement from the 12 available, was selected and the sequences for those proteins were acquired from the 4 selected individuals. Ortholog sequences between the 4 individuals were aligned using Mafft, generating an $N$ number of alignments (one for each protein). The $N$ protein alignments were then concatenated, merging them into a single alignment. The concatenated alignment was then used as input for PhyML, generating a phylogenetic tree for these 4 individuals and the populations they represent. The generated tree was first rooted at a pre-selected outgroup, *Pongo abelii* for the hominid quadruplet and *Pan troglodytes* for the hominin quadruplet. The generated tree was also trimmed for extremely short branches (length <= 0.0000001), which were flattened into polytomies.The topology of the generated tree is then compared to a model tree. The model trees for each quadruplet are showcased in Main text figures 1,5 and 6. Each generated tree is then assigned a label, depending on the topology it presents. This analysis was repeated 1000 times for each $N$, with $N$ scaling from 1 to 12.

##### 3.2 Reproducing the results - Effect of number of proteins in tree inference

All the necessary code and data to reproduce this analysis is available at Github:

`https://github.com/johnpatramanis/Protein_ILS_Hominids_and_Hominins/tree/main`

and require only a Linux machine with conda installed. To reproduce the code however, the user should first install 2 conda environments that make all the tools available. The two environments are available in the main folder and can be installed with the following commands:

```
##### Install Environments
conda env create -f Analyser.yml
conda env create -f Entropy.yml
```

The conda environments can alternatively be created manually with the following commands:

```
#### Install Environment
conda create -n Analyser -c bioconda -c conda-forge snakemake phyml mafft mrbayes revbayes trimal
    bioconductor-shortread r-stringr r-data.table r-phyclust seqmagick

conda create -n Entropy -c conda-forge -c bioconda biopython r-bio3d snakemake biopython
```

Once the environments are installed (or if they have been previously installed), the user can enter the `ILS - Hominids/` folder and type the following:

```
#### Run workflow
bash Master_Script.sh
```

This will generate the results for the hominid itterative analysis. To do the same for the hominin dataset, simply move to the `ILS - Hominins/` folder and type the same:

```
#### Run workflow
bash Master_Script.sh
```

The results were plotted using Python 3 and Matplot lib, by moving into either the `ILS - Hominids/Plotting/` or the `ILS - Hominins/Plotting/` folder and typing the following:

```
#### Run plotting
python3 Distance_Number_Of_Genes.py
```

##### 3.3 Variants Tables

| Number of Proteins | Mean number of variants | Median number of variants | Minimum number of variants | Maximum number of variants |
|---|---|---|---|---|
| 1 | 9.107 | 7.0 | 0 | 36 |
| 2 | 18.721 | 16.0 | 2 | 53 |
| 3 | 28.113 | 26.0 | 6 | 70 |
| 4 | 37.794 | 37.0 | 9 | 73 |
| 5 | 46.971 | 46.5 | 14 | 85 |
| 6 | 57.039 | 58.0 | 23 | 93 |
| 7 | 66.475 | 67.0 | 30 | 103 |
| 8 | 75.125 | 76.0 | 39 | 118 |
| 9 | 84.824 | 87.0 | 44 | 114 |
| 10 | 94.153 | 96.0 | 57 | 121 |
| 11 | 104.013 | 106.0 | 76 | 129 |
| 12 | 113.399 | 113.0 | 102 | 132 |

Table 2: Table containing the mean, median, maximum and minimum number of variants for 12000 replications of the “hominid” dataset’s iterative analysis. Each row corresponds to a number of proteins ranging from 1 to 12. Here a variant is considered a position that has at least one different aminoacid in the alignment of humans, *Pan* and *Gorilla*

| Number of Proteins | Mean number of variants | Median number of variants | Minimum number of variants | Maximum number of variants |
|---|---|---|---|---|
| 1 | 1.0 | 1.0 | 0 | 7 |
| 2 | 2.0 | 2.0 | 0 | 9 |
| 3 | 3.0 | 3.0 | 0 | 10 |
| 4 | 5.0 | 5.0 | 0 | 12 |
| 5 | 6.0 | 6.0 | 0 | 15 |
| 6 | 7.0 | 7.0 | 1 | 15 |
| 7 | 8.0 | 8.0 | 0 | 17 |
| 8 | 9.0 | 9.0 | 2 | 20 |
| 9 | 11.0 | 11.0 | 3 | 19 |
| 10 | 12.0 | 12.0 | 4 | 20 |
| 11 | 13.0 | 13.0 | 5 | 22 |
| 12 | 14.0 | 14.0 | 8 | 21 |

Table 3: Table containing the mean, median, maximum and minimum number of variants for 12000 replications of the "hominin" dataset's iterative analysis. Each row corresponds to a number of proteins ranging from 1 to 12. Here a variant is considered a position that has at least one different aminoacid in the alignment of modern humans, Neanderthals and Denisovans

| Number of Proteins | Mean number of variants | Median number of variants | Minimum number of variants | Maximum number of variants |
|---|---|---|---|---|
| 1 | 1.49 | 1.0 | 0 | 9 |
| 2 | 2.937 | 2.0 | 0 | 12 |
| 3 | 4.386 | 4.0 | 0 | 17 |
| 4 | 5.984 | 5.0 | 0 | 18 |
| 5 | 7.246 | 7.0 | 0 | 20 |
| 6 | 8.876 | 8.0 | 0 | 22 |
| 7 | 10.306 | 10.0 | 1 | 25 |
| 8 | 11.798 | 11.0 | 1 | 25 |
| 9 | 13.268 | 13.0 | 2 | 27 |
| 10 | 14.69 | 14.0 | 3 | 29 |
| 11 | 16.359 | 16.0 | 5 | 34 |
| 12 | 17.553 | 17.0 | 4 | 38 |
| 13 | 19.277 | 19.0 | 7 | 35 |
| 14 | 20.622 | 20.0 | 7 | 35 |
| 15 | 22.086 | 22.0 | 6 | 39 |
| 16 | 23.425 | 23.0 | 9 | 38 |
| 17 | 24.985 | 25.0 | 12 | 44 |
| 18 | 26.753 | 27.0 | 13 | 40 |
| 19 | 27.995 | 28.0 | 15 | 43 |
| 20 | 29.435 | 29.0 | 13 | 43 |
| 21 | 30.916 | 31.0 | 17 | 46 |
| 22 | 32.734 | 33.0 | 19 | 46 |
| 23 | 33.872 | 34.0 | 20 | 47 |
| 24 | 35.345 | 36.0 | 20 | 48 |
| 25 | 36.761 | 37.0 | 25 | 50 |
| 26 | 38.268 | 38.0 | 26 | 51 |
| 27 | 39.628 | 40.0 | 28 | 51 |
| 28 | 41.166 | 41.0 | 32 | 53 |

Table 4: Table containing the mean, median, maximum and minimum number of variants for 28000 replications of the "bone-dentin" dataset's iterative analysis. Each row corresponds to a number of proteins ranging from 1 to 28. Here a variant is considered a position that has at least one different aminoacid in the alignment of modern humans, Neanderthals and Denisovans

##### 3.4 Removal of Collagen

We further investigated whether the discrepancy in the amount of phylogenetic information between the enamel and collagen type I proteins can influence the generated tree topologies for the hominid dataset. To test this, we repeated the hominid tree analysis, this time excluding the 2 collagen type I proteins and using only the enamel specific ones. To reproduce this analysis, simply repeat the steps described in 3.2 with a modification. Simply enter the `ILS - Hominids/` folder and edit the file named ‘‘`Protein_Subset`" by removing the two rows of `COL1A1` and `COL1A2`. Then, repeat the commands as described before in S3.2. Note that this will overwrite any previous results.

Below you can find the two figures generate showcasing the results (see figure 3 and figure 4).

When doing so, we notice a slight increase in the trees in agreement with the known species tree, compared to the full enamel and collagen dataset. While no difference is visible when using between 1 and 5 proteins, there is a noticeable reduction in the number of discordant trees when using between 6 and 8 proteins (or an increase in the trees supporting topology #1).

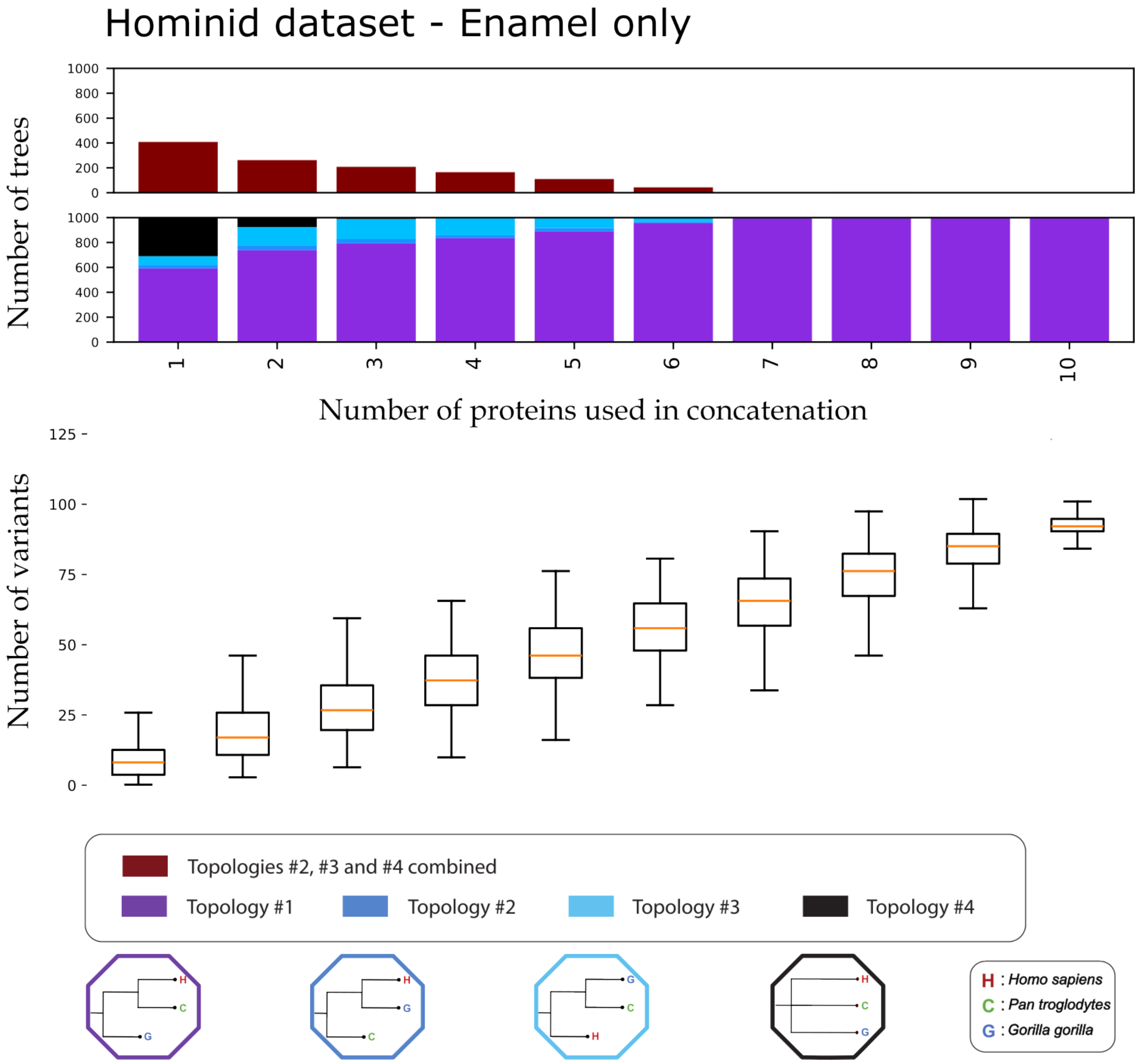


Figure 3: Iterative concatenation analysis for the hominid dataset, but using Enamel proteins only. The upper barplot showcases the number of trees, out of 1000, differing from Topology #1 for an $N$ number of proteins used in the concatenation. The lower barplot showcases the percentage of trees supporting each of the 4 possible topologies, out of 1000, for an $N$ (1-10) number of proteins used in the concatenation. The 4 possible topologies and their corresponding colours are visible at the bottom of the plot. The number of variants present in the dataset creating each tree are visible below the bar plot as a box plot. For each box the orange line denotes the median of the variants present in the $N$ number of proteins concatenation. Each box plot denotes the 25%, 50% (the median, the line in the middle of the box) and 75% quantiles of the distribution. The whiskers of each box denote extremely low values (25% quantile - 1.5 * interquantile range) and extremely high values (75% quantile + 1.5 * interquantile range) for that distribution. The table containing the exact mean, median, maximum and minimum variants for each $N$ proteins is available in Supplementary.

### Comparison of Hominid dataset using only enamel proteins

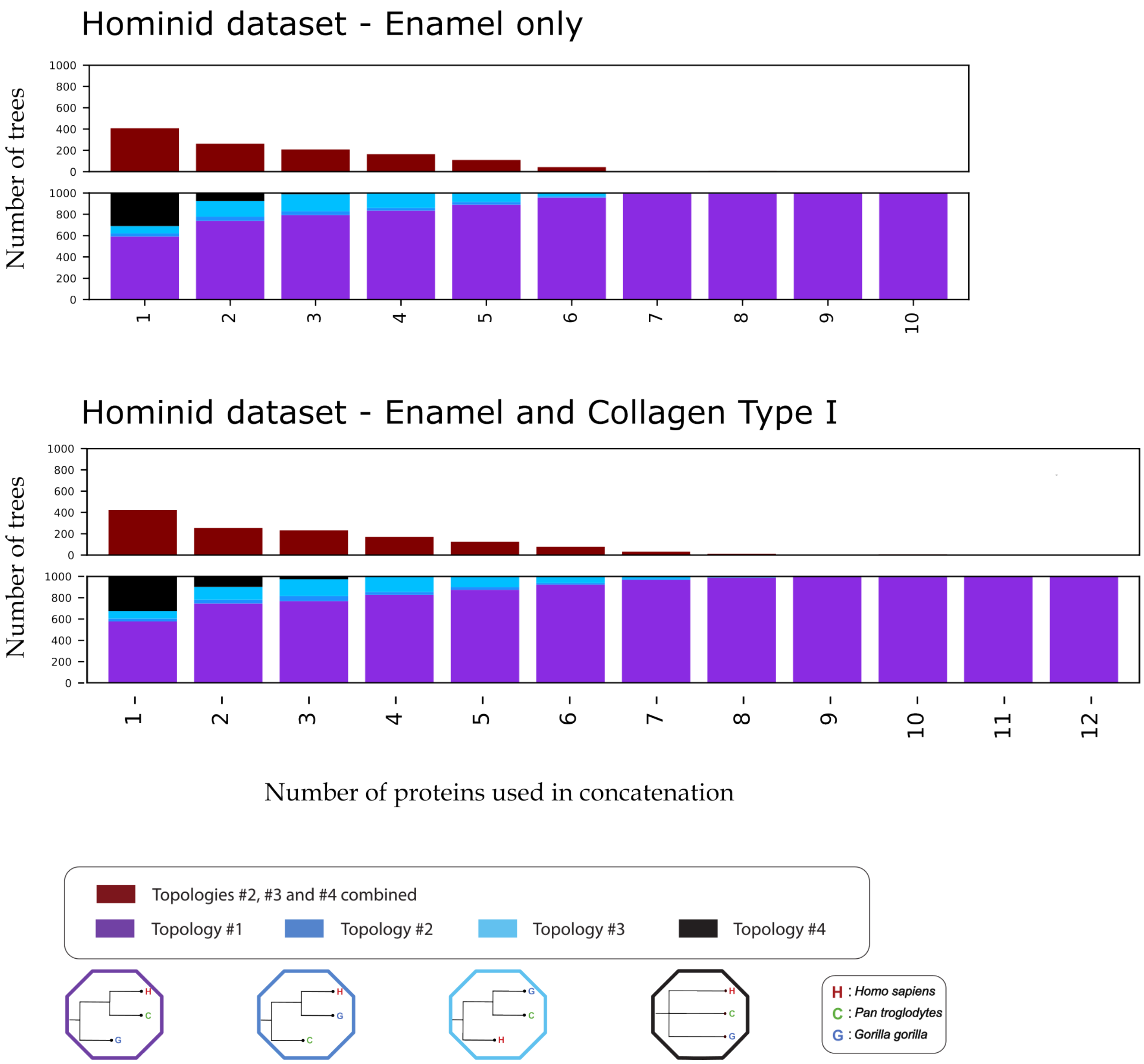


Figure 4: Comparison of results from the iterative analysis of the hominid dataset, using only enamel proteins (top) and both enamel and collagen type I proteins (bottom). For each of the two datasets, the first barplot at the top of the figure represents the number of trees, out of 1000 repetitions, differing from topology , for each number of proteins used in a concatenation, ranging from one to twelve (or one to ten). The second barplots breaks down the number of trees supporting each of the four topologies for the same number of proteins.

##### 3.5 Bootstrap Support of Trees

As described in the methodology section of the main text, we extracted and plotted the bootstrap support of each tree from all iterative analysis datasets. We grouped the bootstrap support scores according to the data set, the number of proteins used to generate them, and the tree topology they supported. We then plotted them as boxplots using python's 3 Matplotlib [13] package, selecting the option to not plot outliers for visual clarity. Below you can find the resulting figures along with a more detailed discussion.

In the hominid data set, we observe that the median bootstrap support of topology #1 steadily increases, from a median of 75% ($N = 1$) to more than 90% ($N>8$), as we increase the number of proteins figure 5. The median support of topology #2, while initially very similar to the support for topology #1, rapidly drops with the addition of more protein sequences. Topology #3, already being weakly supported when using a few proteins, drops even further in support with the addition of more proteins.

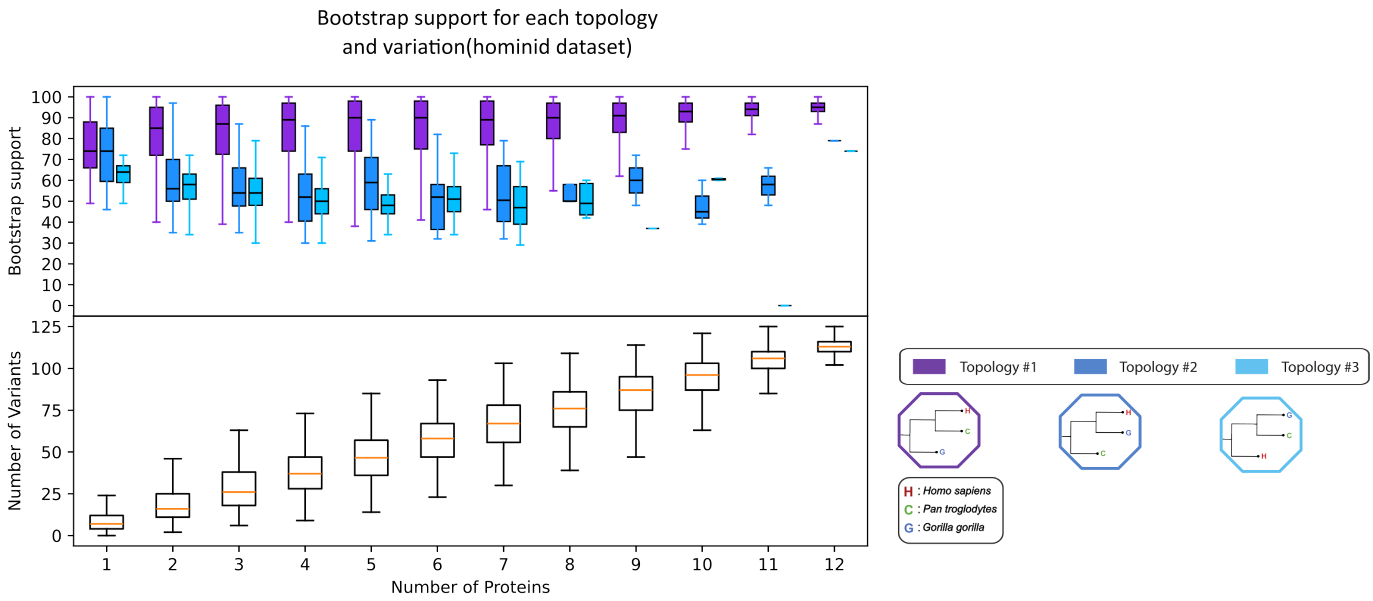


Figure 5: Box plot of the bootstrap values of the hominid dataset's generated trees. A bootstrap value is a metric generated along with a phylogenetic tree, representing the "confidence" of that tree. In our results, the bootstrap values of a tree range between 0 (lowest confidence) to 100 (highest confidence). Each box in this plot represents the distribution of these bootstrap values, grouped together by the number of proteins used to generate their corresponding tree (position on the x axis) and the topology that tree supported (colour). The three topologies corresponding to each colour are shown in the bottom right of the plot. Each box plot denotes the 25%, 50% (the median, the line in the middle of the box) and 75% quantiles of the distribution. The whiskers of each box denote extremely low values (25% quantile - 1.5 * interquantile range) and extremely high values (75% quantile + 1.5 * interquantile range) for that distribution. Each box in the boxplot below shows the number of phylogenetically informative sites (as a distribution) that were used to generate the trees that the bootstrap values originate from.

For the hominin dataset, although a few trees supported topologies #1 and #3 when using a single protein, their median bootstrap support is 0, while topology #2 presented a median of 75% figure 6. This median support for topology #2 slightly drops as the number of proteins increases, but remains roughly the same (support is 69% when using all 12 proteins). Topology #3 shows very low bootstrap support (<60%), dropping further when higher numbers of proteins were used. Finally topology #1, the topology in agreement with the population tree, although initially unsupported, displayed similar bootstrap supports as topology #2 when using more than 3 proteins.

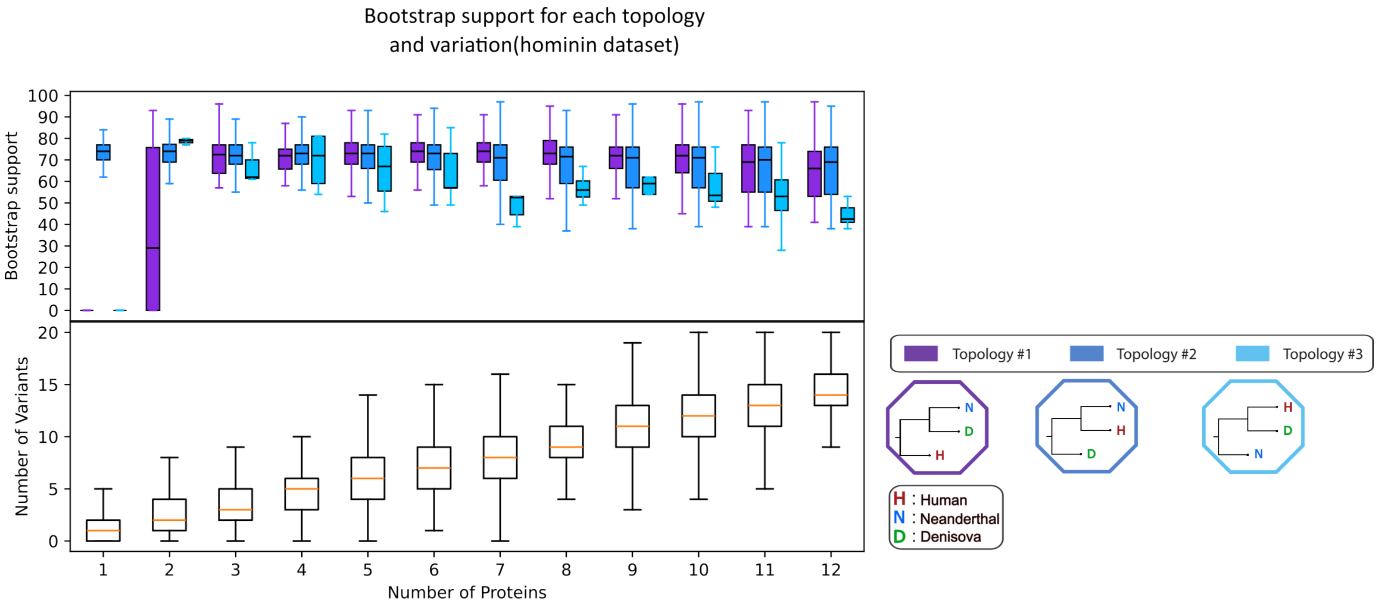


Figure 6: Box plot of bootstrap values of the hominin dataset's generated trees. A bootstrap value is a metric generated along with a phylogenetic tree, representing the "confidence" of that tree. In our results, the bootstrap values of a tree range between 0 (lowest confidence) to 100 (highest confidence). Each box in this plot represents the distribution of these bootstrap values, grouped together by the number of proteins used to generate their corresponding tree (position on the x axis) and the topology that tree supported (colour). The three topologies corresponding to each colour are shown in the bottom right of the plot. Each box plot denotes the 25%, 50% (the median, the line in the middle of the box) and 75% quantiles of the distribution. The whiskers of each box denote extremely low values (25% quantile - 1.5 * interquantile range) and extremely high values (75% quantile + 1.5 * interquantile range) for that distribution. Each box in the boxplot below shows the number of phylogenetically informative sites (as a distribution) that were used to generate the trees that the bootstrap values originate from.

Although the bone-dentin dataset utilized a larger number of available proteins ($n = 28$) the resulting bootstrap supports do not look very different from the hominin dataset. The bootstrap supports of each topology remained roughly stagnant throughout the increase of proteins: Topologies #1 and #2 show almost identical distributions of values, with a fluctuating median of 70%, while topology #3 varies between 45% and 55% figure 7.

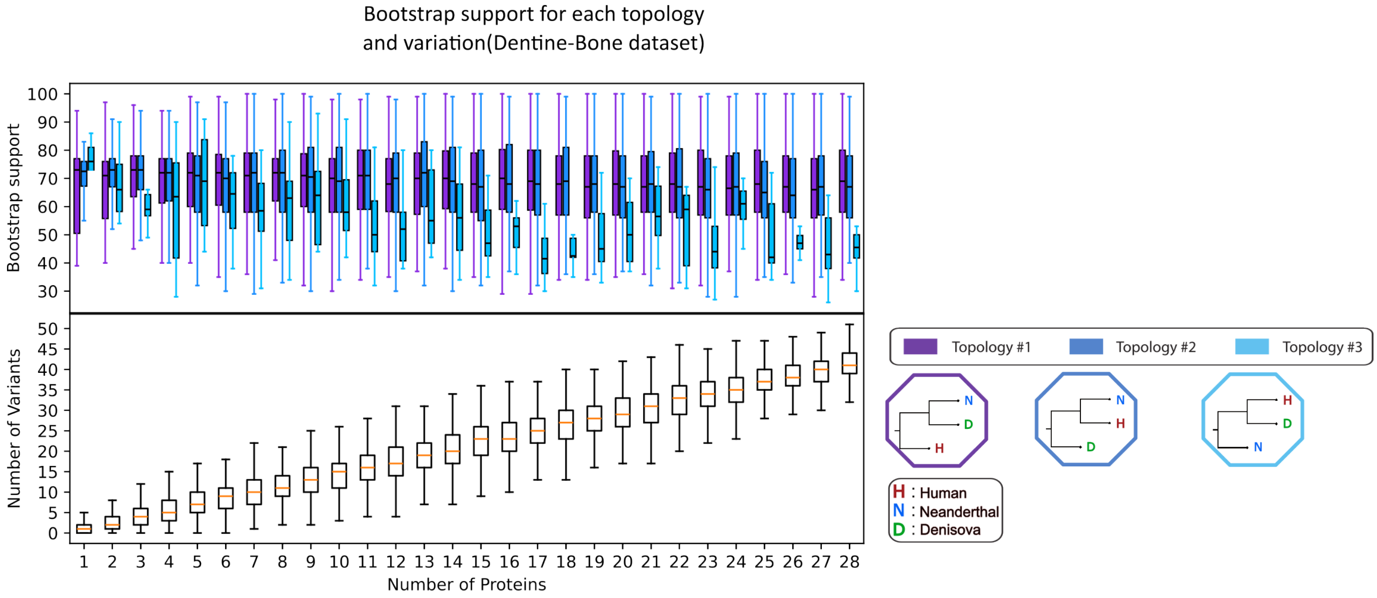


Figure 7: Box plot of bootstrap values of the bone-dentin dataset's generated trees. A bootstrap value is a metric generated along with a phylogenetic tree, representing the "confidence" of that tree. In our results, the bootstrap values of a tree range between 0 (lowest confidence) to 100 (highest confidence). Each box in this plot represents the distribution of these bootstrap values, grouped together by the number of proteins used to generate their corresponding tree (position on the x axis) and the topology that tree supported (colour). The three topologies corresponding to each colour are shown in the bottom right of the plot. Each box plot denotes the 25%, 50% (the median, the line in the middle of the box) and 75% quantiles of the distribution. The whiskers of each box denote extremely low values (25% quantile - 1.5 * interquantile range) and extremely high values (75% quantile + 1.5 * interquantile range) for that distribution. Each box in the boxplot below shows the number of phylogenetically informative sites (as a distribution) that were used to generate the trees that the bootstrap values originate from.

For the repeat analysis of the hominid data set , without the inclusion of Collagen Type I, we observe very similar results to those of the hominid dataset that includes all 12 proteins (see figure figure 8). However, a noticeable difference is the quicker convergence of topology #1 to a high average bootstrap support, coinciding with the rapid drop of the supports for topologies #2 and #3. The results of the non-collagen dataset of around 8 to 9 proteins, mirror that of the collagen-and-enamel dataset at around 11 to 12 proteins. In summary, the removal of collagen ensures a 'faster' convergence to high bootstrap supports for topology #1 and low bootstrap supports for the remaining topologies.

After repeating the analysis of the hominin dataset, but only sampling individuals from Africa we observe roughly similar results, with one point of difference (see figure figure 9). Specifically we notice the complete collapse of support for topology #3 through the iterative analysis, which in the original hominin dataset maintained low but consistent bootstrap support.

When repeating the analysis of the bone-dentin dataset, but only sampling individuals from Africa, we also observe small differences to that of the original dataset (see figure figure 10). Here we notice a substantial decrease for topology #3, which sees a continual drop along the increase of protein data, all the way to a mean of around

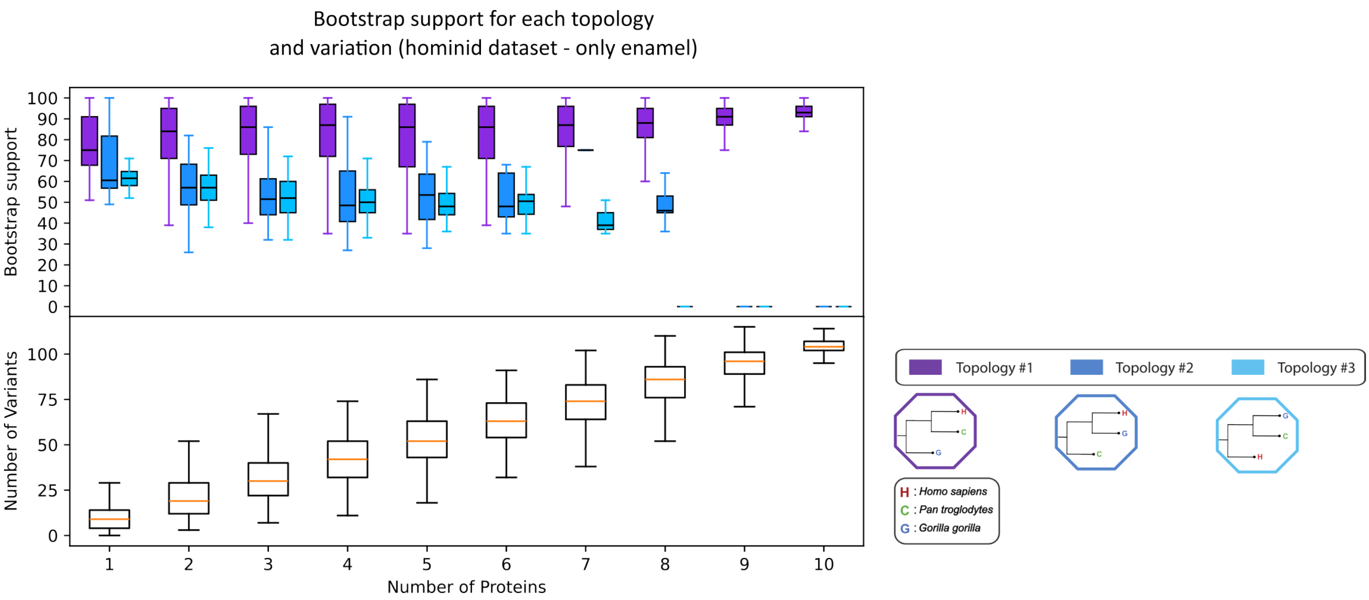


Figure 8: Box plot of bootstrap values of the hominid dataset's generated trees, using only the 10 enamel-specific proteins. A bootstrap value is a metric generated along with a phylogenetic tree, representing the "confidence" of that tree. In our results, the bootstrap values of a tree range between 0 (lowest confidence) to 100 (highest confidence). Each box in this plot represents the distribution of these bootstrap values, grouped together by the number of proteins used to generate their corresponding tree (position on the x axis) and the topology that tree supported (colour). The three topologies corresponding to each colour are shown in the bottom right of the plot. Each box plot denotes the 25%, 50% (the median, the line in the middle of the box) and 75% quantiles of the distribution. The whiskers of each box denote extremely low values (25% quantile - 1.5 * interquantile range) and extremely high values (75% quantile + 1.5 * interquantile range) for that distribution. Each box in the boxplot below shows the number of phylogenetically informative sites (as a distribution) that were used to generate the trees that the bootstrap values originate from.

50 for 28 proteins.

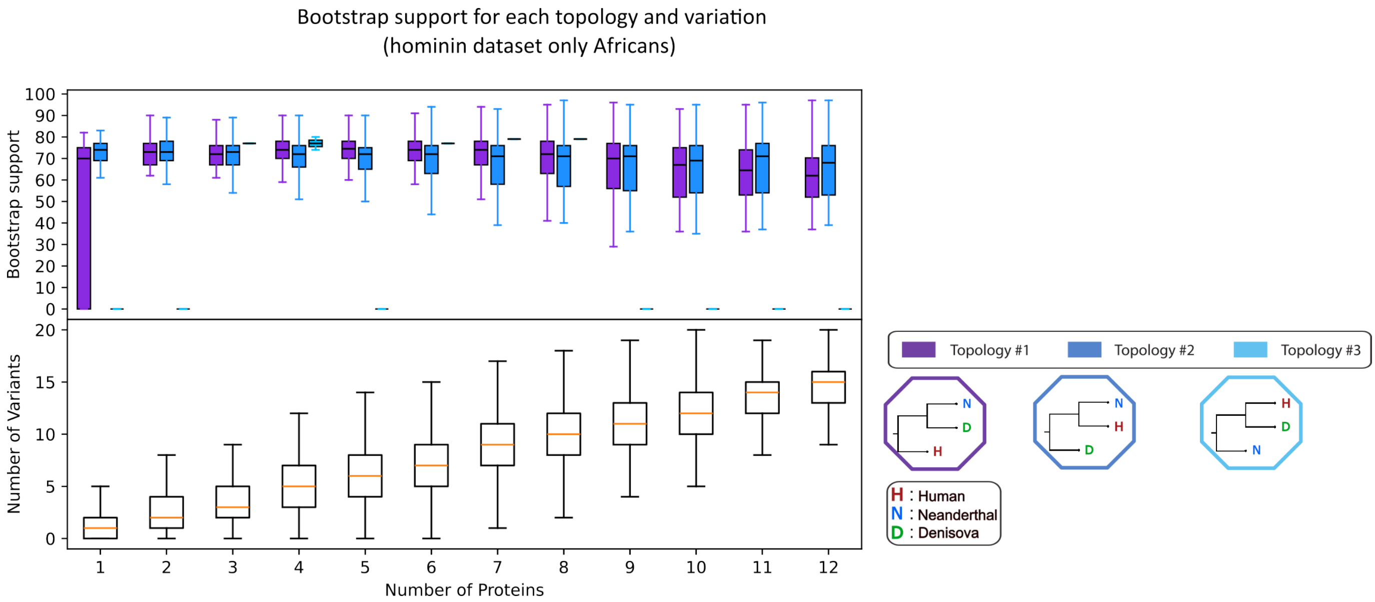


Figure 9: Box plot of bootstrap values of the hominin dataset's generated trees, but using only individuals from specific African populations as humans. A bootstrap value is a metric generated along with a phylogenetic tree, representing the "confidence" of that tree. In our results, the bootstrap values of a tree range between 0 (lowest confidence) to 100 (highest confidence). Each box in this plot represents the distribution of these bootstrap values, grouped together by the number of proteins used to generate their corresponding tree (position on the x axis) and the topology that tree supported (colour). The three topologies corresponding to each colour are shown in the bottom right of the plot. Each box plot denotes the 25%, 50% (the median, the line in the middle of the box) and 75% quantiles of the distribution. The whiskers of each box denote extremely low values (25% quantile - 1.5 * interquantile range) and extremely high values (75% quantile + 1.5 * interquantile range) for that distribution. Each box in the boxplot below shows the number of phylogenetically informative sites (as a distribution) that were used to generate the trees that the bootstrap values originate from.

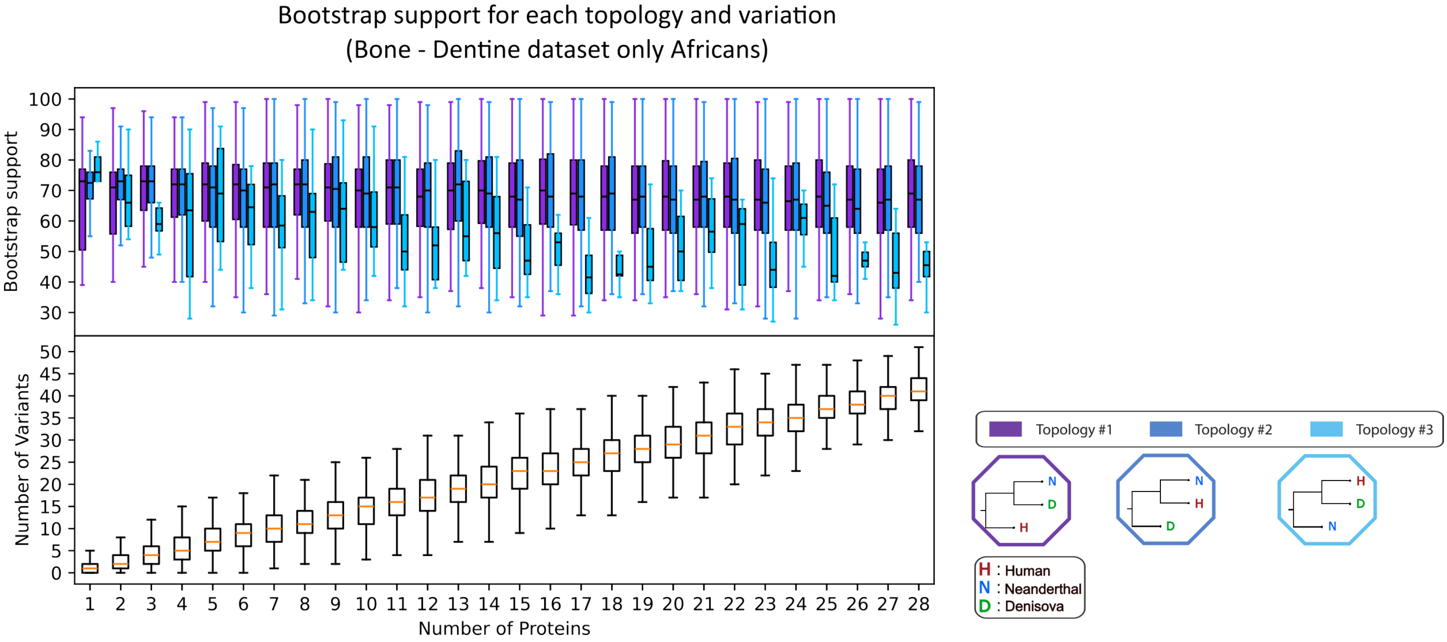


Figure 10: Box plot of bootstrap values of the bone-dentin dataset's generated trees, but using only individuals from specific African populations as humans. A bootstrap value is a metric generated along with a phylogenetic tree, representing the "confidence" of that tree. In our results, the bootstrap values of a tree range between 0 (lowest confidence) to 100 (highest confidence). Each box in this plot represents the distribution of these bootstrap values, grouped together by the number of proteins used to generate their corresponding tree (position on the x axis) and the topology that tree supported (colour). The three topologies corresponding to each colour are shown in the bottom right of the plot. Each box plot denotes the 25%, 50% (the median, the line in the middle of the box) and 75% quantiles of the distribution. The whiskers of each box denote extremely low values (25% quantile - 1.5 * interquantile range) and extremely high values (75% quantile + 1.5 * interquantile range) for that distribution. Each box in the boxplot below shows the number of phylogenetically informative sites (as a distribution) that were used to generate the trees that the bootstrap values originate from.

### 4. Introgression Investigation

##### 4.1 Overlap of Enamel Genes and identified introgressed regions in modern human populations.

We investigated whether the inability to resolve the human - Neanderthal - Denisovan phylogenetic relations, using the proteins of this study, could be a result of past admixture events between the 3 populations. For this, we looked at previously published archaic-introgressed segments of modern humans. These are segments that have been inferred to be present in modern human genomes but originating from either Neanderthals, Denisovans or possibly other archaic humans. We used two lists of genetic coordinates from Skov et al. [14] and Chen et al. [15], pertaining segments that were inferred as being introgressed from archaic groups.

Below we describe in detail the process for identifying if a selection of proteins (the genes coding them) are located within archaic-introgressed segments. We will first use the data produced using Hmmix from from Skov et al. [14] and then repeat this process for IBDmix from Chen et al. [15].

4.1.1 Introgressed proteins based on Hmmix.

All the necessary code and data to reproduce this analysis is available at:

`https://github.com/johnpatramanis/Protein_ILS_Hominids_and_Hominins/tree/main`

and requires only a Linux machine with Python 3 installed to run. To reproduce the analysis, first navigate to the appropriate directory:

```
##### Move to directory
cd Introgression_Investigation/Skov_et_al_2024/
```

Unfortunately the data to reproduce this analysis are a bit large (more than 1.5 Gb of space) and need to be downloaded separately from Zenodo (https://zenodo.org/records/14136628). This can be done manually by pressing the download button next to the file `hg38_1000g_segments.txt` and `hg38_HGDP_segments.txt`. These two files should be placed within the `Introgression_Investigation\Skov_et_al_2018` folder. Alternatively the following command in a linux environment will download them:

```
##### Download Introgression Data from Skov et al
wget https://zenodo.org/records/14136628/files/hg38_1000g_segments.txt
wget https://zenodo.org/records/14136628/files/hg38_HGDP_segments.txt
```

Make sure both files are present inside the `Introgression_Investigation` folder before proceeding. The two lists correspond to two different global datasets of modern humans: the 1000 Genomes Project (1GP) and the Human Genome Diversity Project (HGDP). These two text files are long lists containing information on segments of the human genome that have previously been inferred to be introgressed from other archaic human groups, through the process of admixture. Each row has a single entry of an introgressed segment from a single individual genom. This includes information on the start and end of the segment (in genetic coordinates), the modern human population which the individual belongs to (as it is labeled in the original 1GP or HGDP data), the archaic human genomes that match the intogressed segment as well as a probability that the segment is truly introgressed. Important to note here, both the coordinates of the gene of interest and the data from Skov et al. correspond to GrCh38.

Now we will now run a script that will compare the genetic coordinates of the proteins under investigation and the coordinates of the introgressed segments to identify any potential overlaps. Note that the coordiantes of the genes coding the proteins of interest can be found in `Protein_Location_Sorted.txt` , but the user is free to add or remove proteins from this set. If so, please make sure the format is the same: `GeneName Chromosome Start End`.

```
#### Execute python script
python3 Check_Overlap.py
```

After running, this will generate a new file named `Introgressed_Proteins.txt`. We will now run another short python script, to gather information on each protein and print it in a short report. Run the following command to do so:

```
#### Execute python script
python3 Report_on_found_introgressed_segments.py
```

The information on the proteins found within introgressed segments will now be printed on your screen, and inside the file `Report_on_Introgressed_Proteins.txt`. The results can be seen here in Table 5. We have also extended the above methodology to the Dentin-bone dataset of proteins ($n = 28$) and seperated the results per continental population and by the two archaic populations in Tables 6 and 7.

We can also visualize this overlap between introgressed segments and the proteins of interest, using a combination of a python and R script. We chose to use MMP20 as an example, due to the large amount of introgression that is detected, but this should work with any protein.

```
#### move to plotting directory
cd Plotting

#### download some additional files from Skov et al 2024 Zenodo page
wget https://zenodo.org/records/14136628/files/hg38_1000g_SNPS.txt
wget https://zenodo.org/records/14136628/files/hg38_HGDP_SNPS.txt

##### Exectute python script, giving a name for the gene/protein and its coordinates
python Get_data.py -outfile=MMP20 -chrom=chr11 -start=102_576_832 -end=102_625_332

#### Plot results with R
Rscript plot.R MMP20 chr11 102576832 102625332
```

The above code should have now generated a PDF file named `chr11_102576832_102625332_MMP20.pdf`, which should match Main text figure 7.

##### 4.1.2 Introgressed proteins based on IBDmix.

Here we repeat the identification of possibly introgressed proteins, using the archaic-introgressed haplotypes of modern humans, classified as such by IBDmix. IBDmix, published in Chen et al. [15] has been used to identify introgressed Neanderthal segments in modern humans. Although Chen and colleagues did not include Denisovan introgression in their analysis, their results can still be used to confirm that some of the enamel and collagen proteins under investigation may originate from archaic humans (Neanderthals). We repeated the same process as with the data from Skov and colleagues. An important note is that we first translated the coordinates of the genes under investigation (from GrCh38) to match the reference genome of the data of Chen and colleagues (to GrCh37). Additionally, the analysis of Chen et al. [15] consists of 3 separate runs using 3 different Neanderthal genomes. As a results a number of introgressed segments seen in column 3 of 8 may be duplicates. For column 3 of the same table, only one Neanderthal genome was used to calculate the frequency in modern humans. To repeat this analysis follow the instructions below.

```
##### Move to Chen_et_al_2020 directory from Skov_et_al_2024
cd ../Chen_et_al_2020/
##### Download Introgression Data from Chen et al, unpack it and move folders to directory here
wget
    https://github.com/PrincetonUniversity/IBDmix/archive/refs/heads/main/IBDmix_calls_using_3_archaics.tar.gz
tar -xvzf IBDmix_calls_using_3_archaics.tar.gz
tar -xvzf IBDmix-main/IBDmix_calls_using_3_archaics.tar.gz
mv -f ./IBDmix-main/IBDmix_calls_using_3_archaics/* ./

#### Execute python script
python3 Report_on_found_introgressed_segments.py
#### Execute python script
python3 Report_on_found_introgressed_segments.py
```

Again, the information on the proteins found within introgressed segments will now be printed on your screen, and inside the file `Report_on_Introgressed_Proteins.txt`, but also here in Table 8.

| Protein Name | Number of identified haplotypes | Fraction of introgressed haplotype | Population of the individuals the haplotypes | Matching ancient genomes |
|---|---|---|---|---|
| MMP20 | 1216 | 0.182 | Adygei, Tu, Tuscan, Tujia, STU, MXL, Cambodian, Yakut, Mozabite, Palestinian, Orcadian, Dai, Brahui, Burusho, CDX, FIN, She, NorthernHan, Pima, Makrani, Sindhi, Druze, PJL, CLM, CEU, Oroqen, Balochi, Surui, Karitiana, Naxi, PEL, Hezhen, Pathan, Lahu, KHV, ITU, Russian, Japanese, Yi, Mongolian, Basque, Daur, Bedouin, Kalash, French, Uygur, PUR, Colombian, BergamoItalian, BEB, GBR, CHB, Sardinian, Maya, Xibo, GIH, Han, Miao, CHS, JPT, Hazara, TSI, IBS | Neanderthal, Denisova, none |
| COL1A1 | 23 | 0.003 | Pathan, Lahu, NorthernHan, CHB, Naxi, Han, Yakut, Miao, Palestinian, CHS, JPT, Bedouin, PEL, Hezhen | Neanderthal, none, Denisova |
| COL17A1 | 2 | 0.0003 | KHV | none |
| ODAM | 32 | 0.005 | Tu, ITU, PapuanHighlands, STU, BEB, PJL, GIH, PapuanSepik, Hazara, Bougainville | none, Denisova |
| AMTN | 22 | 0.003 | ITU, STU, BEB, PJL, GIH | Denisova |
| AMBN | 27 | 0.004 | Pathan, ITU, STU, BEB, Makrani, PJL, GIH | Denisova |
| ALB | 31 | 0.005 | Pathan, ITU, PapuanHighlands, STU, BEB, PJL, PapuanSepik, Kalash, Bougainville | Denisova |
| ENAM | 2 | 0.0003 | STU | Denisova |
| COL1A2 | 3 | 0.0004 | Druze, Uygur | none |

Table 5: Results of the introgressed loci investigation using the Skov et al. 2018 results, on the 1000 Genomes and HGDP data. The first column contains the name of the gene investigated. The second column contains the number of introgressed haplotypes found overlapping partially or fully with that specific gene. The third column contains the fraction of these haplotypes that appear within present-day humans of the dataset (globally). The fourth column contains the population names, as they appear in the 1000 Genomes and HGDP dataset, of the modern population the introgressed haplotype was found in. The last column contains the names of archaic genomes that match the allelic state of the introgressed haplotype.

| Gene | Length of Gene (bp) | America | South Asia | East Asia | Europe | Oceania | Dataset (Enamel and Collagen or extended Dentin and Bone) |
|---|---|---|---|---|---|---|---|
| COL11A1 | 232050 | 0 | 0 | 0 | 0 | 0 | Dentin and Bone |
| COL3A1 | 38374 | 0.45 | 3.63 | 3.16 | 3.93 | 0 | Dentin and Bone |
| COL5A2 | 147864 | 0 | 0.13 | 1.24 | 0.19 | 0 | Dentin and Bone |
| COL4A4 | 161492 | 0.82 | 1.07 | 0.37 | 2.41 | 0 | Dentin and Bone |
| AHSG | 8259 | 0 | 0 | 0 | 0 | 0 | Enamel and Collagen |
| ODAM | 8852 | 0 | 0 | 0 | 0 | 0 | Enamel and Collagen |
| AMTN | 14175 | 0 | 0 | 0 | 0 | 0 | Enamel and Collagen |
| AMBN | 15033 | 0 | 0 | 0 | 0 | 0 | Enamel and Collagen |
| ENAM | 18081 | 0 | 0 | 0 | 0 | 0 | Enamel and Collagen |
| ALB | 17196 | 0 | 0 | 0 | 0 | 0 | Enamel and Collagen |
| COL11A2 | 29774 | 29.76 | 13.91 | 6.87 | 12.18 | 1.79 | Dentin and Bone |
| COL12A1 | 121728 | 9.07 | 12.28 | 42.7 | 2.09 | 7.14 | Dentin and Bone |
| COL1A2 | 36333 | 0 | 0 | 0 | 0 | 0 | Enamel and Collagen |
| COL22A1 | 325807 | 3.54 | 3.07 | 0.06 | 7.36 | 0 | Dentin and Bone |
| COL5A1 | 203041 | 8.08 | 33.58 | 1.55 | 15.04 | 37.5 | Dentin and Bone |
| COL17A1 | 54595 | 0 | 0 | 0 | 0 | 0 | Enamel and Collagen |
| F2 | 20294 | 0 | 0 | 0 | 0 | 0 | Dentin and Bone |
| MMP20 | 48501 | 36.84 | 29.39 | 39.54 | 41.75 | 16.07 | Enamel and Collagen |
| **COL2A1** | **31510** | **0** | **1.19** | **0.06** | **0** | **0** | **Enamel and Collagen** |
| LUM | 8866 | 0 | 0 | 0 | 0 | 1.79 | Dentin and Bone |
| POSTN | 36184 | 0 | 0 | 0 | 0 | 0 | Dentin and Bone |
| SERPINF1 | 15506 | 0.27 | 0.69 | 0 | 0.57 | 0 | Dentin and Bone |
| COL1A1 | 17531 | 0 | 0 | 0 | 0 | 0 | Enamel and Collagen |
| CHAD | 4385 | 0 | 0 | 0 | 0 | 0 | Dentin and Bone |
| COL5A3 | 50944 | 12.07 | 12.16 | 4.64 | 4.25 | 0 | Dentin and Bone |
| AMELX | 7349 | - | - | - | - | - | Enamel and Collagen |
| BGN | 14567 | - | - | - | - | - | Dentin and Bone |

Table 6: Frequencies of the Neanderthal-assigned introgressed loci from Skov et al. 2018 results, on the 1000 Genomes and HGDP data. Columns 3 to 7 show the frequency of the archaic haplotype for each continent-population as a percentage (e.g. 1.2 = 0.012 = one point two percent). For an exact list of which population belongs to which continent population, see the original data at https://zenodo.org/records/14136628. The last column contains the name of the dataset the protein belongs to, either "Enamel and Collagen" or "Dentin and Bone". Note that AMELX and BGN could not be assessed because the X chromosome is missing from the original analysis

| Gene | Length of Gene (bp) | America | South Asia | East Asia | Europe | Oceania | Dataset (Enamel and Collagen or extended Dentin and Bone) |
|---|---|---|---|---|---|---|---|
| COL11A1 | 232050 | 0 | 0 | 0 | 0 | 1.79 | Dentin and Bone |
| COL3A1 | 38374 | 0.09 | 0 | 0 | 0 | 0 | Dentin and Bone |
| COL5A2 | 147864 | 0.18 | 0 | 0 | 0.06 | 0 | Dentin and Bone |
| COL4A4 | 161492 | 18.06 | 11.78 | 0.19 | 13.26 | 0 | Dentin and Bone |
| AHSG | 8259 | 0 | 0 | 0 | 0 | 7.14 | Enamel and Collagen |
| ODAM | 8852 | 0 | 1.82 | 0 | 0 | 26.79 | Enamel and Collagen |
| AMTN | 14175 | 0 | 1.94 | 0 | 0 | 0 | Enamel and Collagen |
| AMBN | 15033 | 0 | 2.19 | 0 | 0 | 0 | Enamel and Collagen |
| ENAM | 18081 | 0 | 2.19 | 0 | 0 | 0 | Enamel and Collagen |
| ALB | 17196 | 0 | 1 | 0 | 0 | 26.79 | Enamel and Collagen |
| COL11A2 | 29774 | 0 | 0.38 | 0 | 0 | 0 | Dentin and Bone |
| COL12A1 | 121728 | 0 | 0 | 0 | 0 | 17.86 | Dentin and Bone |
| COL1A2 | 36333 | 0 | 0 | 0 | 0 | 0 | Enamel and Collagen |
| COL22A1 | 325807 | 0 | 1.69 | 0 | 0 | 28.57 | Dentin and Bone |
| COL5A1 | 203041 | 0 | 0.13 | 0.87 | 0.13 | 0 | Dentin and Bone |
| COL17A1 | 54595 | 0 | 0 | 0 | 0 | 0 | Enamel and Collagen |
| F2 | 20294 | 0 | 0 | 0 | 0 | 0 | Dentin and Bone |
| MMP20 | 48501 | 0 | 0.81 | 0 | 0.13 | 0 | Enamel and Collagen |
| COL2A1 | 31510 | 0 | 0 | 0 | 0 | 0 | **Enamel and Collagen** |
| LUM | 8866 | 0 | 0 | 0 | 0 | 21.43 | Dentin and Bone |
| POSTN | 36184 | 0 | 0 | 0.12 | 0 | 0 | Dentin and Bone |
| SERPINF1 | 15506 | 0 | 0 | 0 | 0 | 12.5 | Dentin and Bone |
| COL1A1 | 17531 | 0.18 | 0 | 1.24 | 0 | 0 | Enamel and Collagen |
| CHAD | 4385 | 0 | 0 | 0 | 0 | 0 | Dentin and Bone |
| COL5A3 | 50944 | 0 | 0 | 0 | 0 | 21.43 | Dentin and Bone |
| AMELX | 7349 | - | - | - | - | - | Enamel and Collagen |
| BGN | 14567 | - | - | - | - | - | Dentin and Bone |

Table 7: Frequencies of the Denisovan-assigned introgressed loci from Skov et al. 2018 results, on the 1000 Genomes and HGDP data. Columns 3 to 7 show the frequency of the archaic haplotype for each continent-population as a percentage (e.g. 1.2 = 0.012 = one point two percent). For an exact list of which population belongs to which continent population, see the original data at https://zenodo.org/records/14136628. The last column contains the name of the dataset the protein belongs to, either "Enamel and Collagen" or "Dentin and Bone". Note that AMELX and BGN could not be assessed because the X chromosome is missing from the original analysis.

| Protein Name | Number of individuals with haplotype* | Frequency of introgressed haplotype** | Population of the individuals | Found in the Neanderthals genomes |
|---|---|---|---|---|
| MMP20 | 1144 | 0.158 | CHB, IBS, ITU, ASW, PUR, CLM, KHV, PEL, GBR, FIN, CDX, BEB, JPT, ACB, CHS, MXL, GIH, TSI, CEU, STU, PJL | Altai, Vindija, Chagyrskaya |
| COL17A1 | 1 | 0.0003 | PEL | Chagyrskaya |

Table 8: Results of the introgressed loci investigation using the Chen et al. 2020 results, on the 1000 Genomes dataset. The first column contains the name of the gene investigated. The second column contains the number of introgressed haplotypes found overlapping partially or fully with that specific gene. This number differs from that of table 1, as each of the 3 Neanderthal genomes was used in a separate analysis and thus the same haplotype from a modern human can give a positive result of introgression for each one of them. This can lead to the same haplotype being counted twice or thrice. The third column contains the fraction of these haplotypes that appear within present-day humans of the dataset (globally). The fourth column contains the population names, as they appear in the 1000 Genomes, of the modern population the introgressed haplotype was found in. The last column contains the names of archaic genomes that were used to identify the introgressed haplotype.

##### 4.2 Iterative Analysis using only humans from African populations.

As described in the main text, we repeated the iterative analysis for the hominin and dental-bone datasets, to assess the effect of using un-admixxed individuals when generating the phylogenetic trees. For this, we selected only modern humans from the 1000 genomes populations of Yoruba, Mende, Luhya and Mandinka, as the modern human representative, repeating the analysis. Below you can see the full results of this analysis for the hominin and dental-bone dataset (figures 11 and 12).

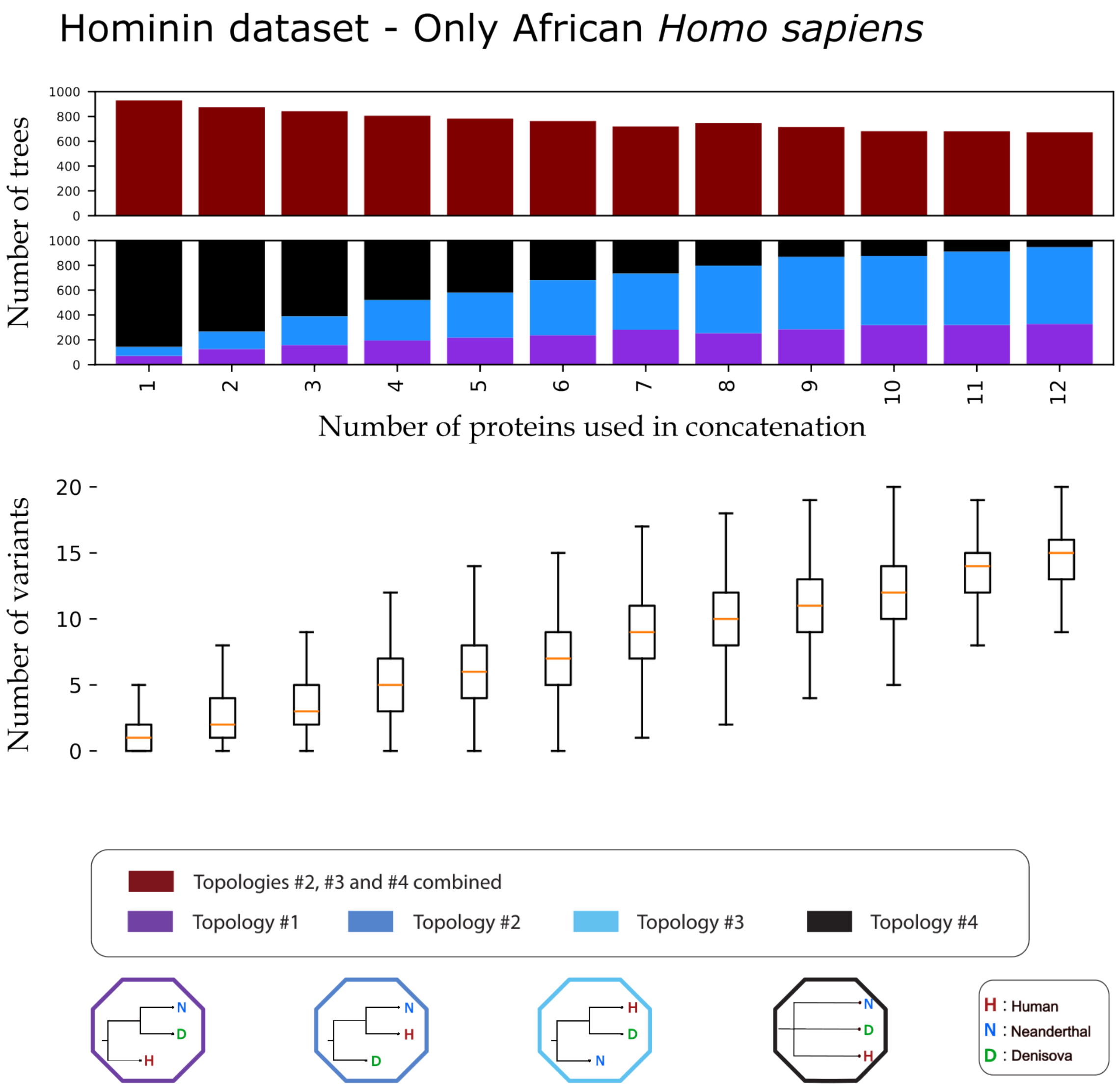


Figure 11: Iterative concatenation analysis for the hominin dataset, but sampling only modern humans from the Yoruba, Mende, Luhya and Mandinka populations. The upper barplot showcases the number of trees, out of 1000, differing from Topology #1 for an $N$ number of proteins used in the concatenation. The lower barplot showcases the percentage of trees supporting each of the 4 possible topologies, out of 1000, for an $N$ (1-10) number of proteins used in the concatenation. The 4 possible topologies and their corresponding colours are visible at the bottom of the plot. The number of variants present in the dataset creating each tree are visible below the bar plot as a box plot. For each box the orange line denotes the median of the variants present in the $N$ number of proteins concatenation. Each box plot denotes the 25%, 50% (the median, the line in the middle of the box) and 75% quantiles of the distribution. The whiskers of each box denote extremely low values (25% quantile - 1.5 * interquantile range) and extremely high values (75% quantile + 1.5 * interquantile range) for that distribution. The table containing the exact mean, median, maximum and minimum variants for each $N$ proteins is available in Supplementary.

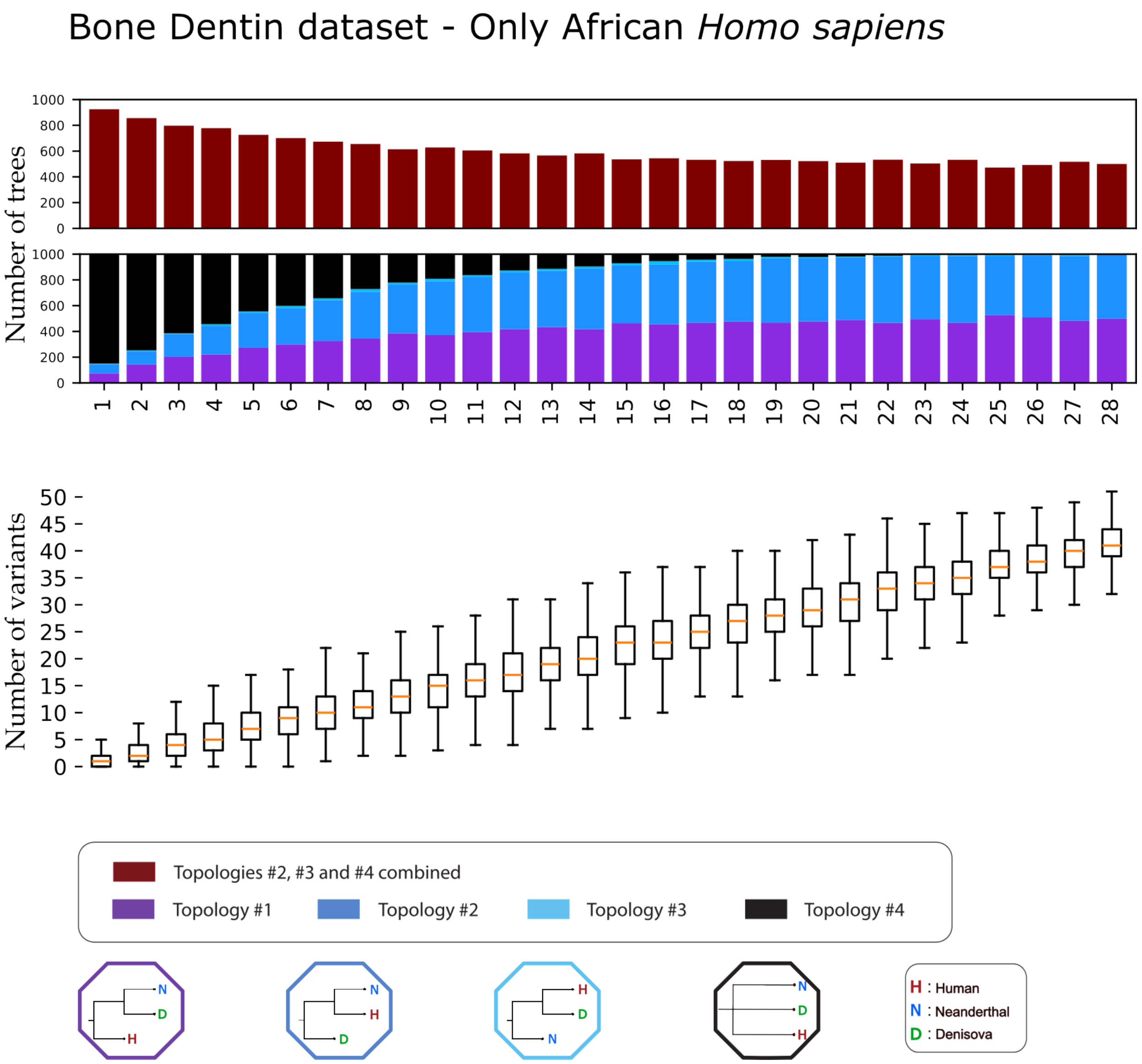


Figure 12: Iterative concatenation analysis for the Dentin-bone dataset, but sampling only modern humans from the Yoruba, Mende, Luhya and Mandinka populations. The upper barplot showcases the number of trees, out of 1000, differing from Topology #1 for an $N$ number of proteins used in the concatenation. The lower barplot showcases the percentage of trees supporting each of the 4 possible topologies, out of 1000, for an $N$ (1-10) number of proteins used in the concatenation. The 4 possible topologies and their corresponding colours are visible at the bottom of the plot. The number of variants present in the dataset creating each tree are visible below the bar plot as a box plot. For each box the orange line denotes the median of the variants present in the $N$ number of proteins concatenation. Each box plot denotes the 25%, 50% (the median, the line in the middle of the box) and 75% quantiles of the distribution. The whiskers of each box denote extremely low values (25% quantile - 1.5 * interquantile range) and extremely high values (75% quantile + 1.5 * interquantile range) for that distribution. The table containing the exact mean, median, maximum and minimum variants for each $N$ proteins is available in Supplementary.

### Comparison of Bone Dentin dataset using only African *Homo sapiens* samples

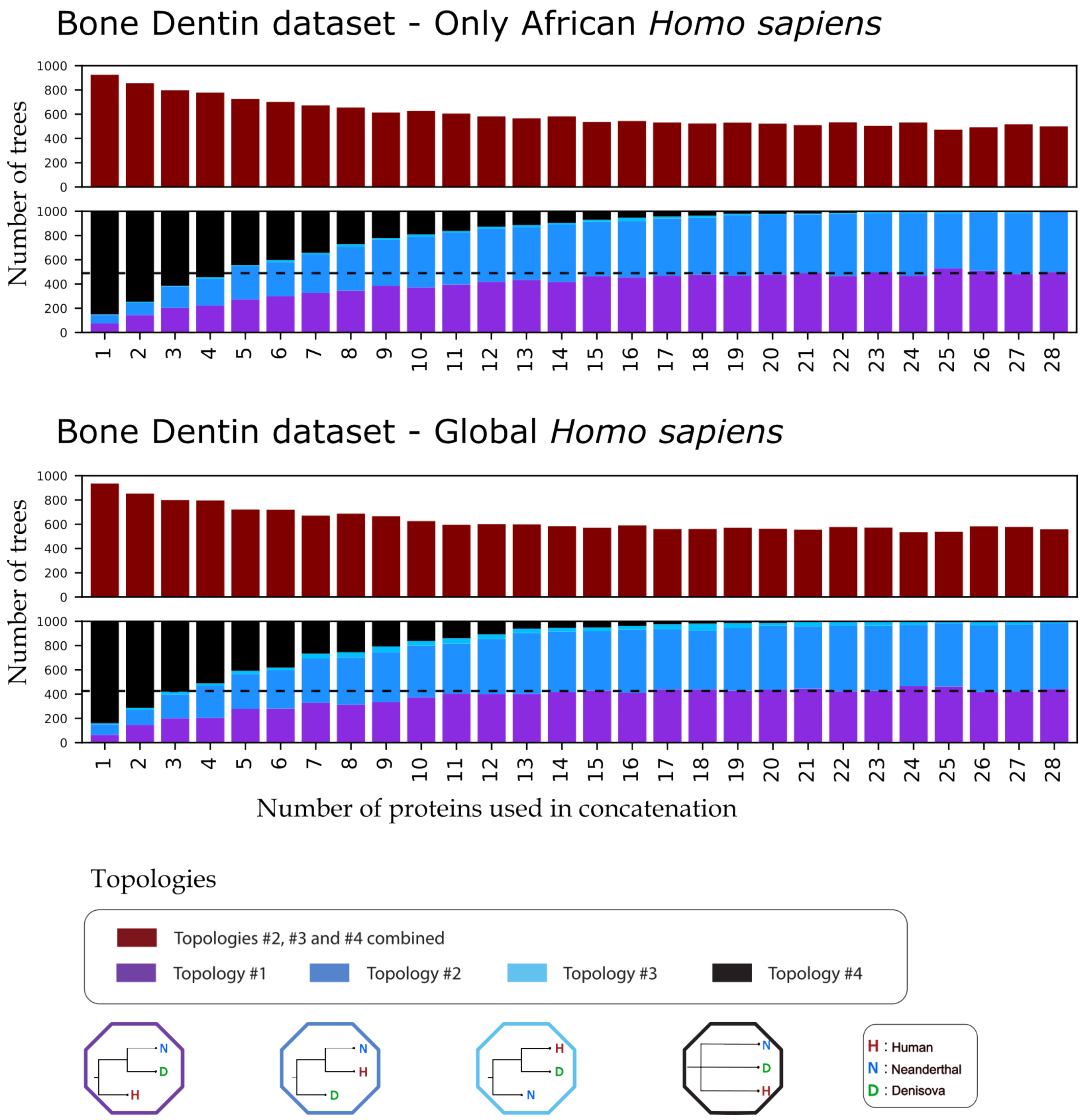


Figure 13: Repeated iterative analysis of the bone-dentin dataset, sampling only individuals belonging to the four populations of Yoruba, Mende, Luhya and Mandinka as "Humans". The first barplot at the top of the figure represents the number of trees, out of 1000 repetitions, differing from topology 1#, for each number of proteins used in a concatenation, ranging from one to twelve. The second barplots breaks down the number of trees supporting each of the four topologies for the same number of proteins. The boxplot below the two barplots represents the number of informative variants in the concatenated alignment of that number of proteins, each number being a distribution of 1000 repetitions.
